# Spatially varying mRNA decay contributes to sharpening the *even-skipped* expression pattern in the *Drosophila* embryo

**DOI:** 10.1101/2025.02.08.637258

**Authors:** Logan Brase, Alastair Pizzey, Jennifer C. Love, Catherine Sutcliffe, Lauren Forbes Beadle, Magnus Rattray, Hilary L. Ashe

**Affiliations:** Faculty of Biology, Medicine and Health, University of Manchester, Manchester, M13 9PT, UK

## Abstract

The regulation of mRNA decay is important for numerous cellular and developmental processes. Here, we use the patterning gene *even-skipped* (*eve*) in the early *Drosophila* embryo to investigate the contribution of mRNA decay to shaping the mature striped expression pattern. Using mathematical models to analyse live and fixed imaging data, we show that for *eve* stripes 2 and 4, spatially regulated degradation rates outperform models with constant or mRNA age-dependent decay. Uniform mRNA decay rates perturb definition of *eve* stripes 2 and 4 *in silico*, and we show that altering *eve* mRNA numbers *in vivo* by changing the UTR results in dysregulation of the anterior-posterior pair-rule network and embryo segmentation defects. Overall, these data demonstrate how *eve* mRNA instability can function with transcriptional regulation to define sharp expression domain borders. We suggest that spatially regulated mRNA stability may be widely used to sculpt expression patterns during development.

## Introduction

mRNA abundance is shaped by a balance of production and degradation. Different combinations of transcription, splicing and degradation rates have been shown to produce similar mRNA expression profiles in mammalian cells, indicating that studies of the regulation of gene expression require an understanding of all parameters.^1,2^ Studying transcription at single cell resolution has revealed how transcriptional parameters vary by cellular position to modulate the mRNA output and therefore downstream processes.^3,4^ Compared to the mechanistic understanding of transcription, much less is understood about how gene expression outcomes are regulated post-transcriptionally.

Spatiotemporal mRNA degradation has been studied in zebrafish and *C. elegans* embryogenesis through a combination of single-cell sequencing and metabolic labeling.^5,6^ The resulting whole-embryo atlases have uncovered different mRNA decay rates across cell types and embryonic stages, indicating that mRNA stability is a highly regulated gene expression step. Estimation of zygotic mRNA half-lives in the early *Drosophila* embryo has revealed that a range of mature mRNA dynamics can arise from similar transcriptional profiles through the regulation of mRNA stability.^7^ Additionally, simulations suggest that mRNA decay contributes to the dynamic shifts in the gap gene expression patterns in the *Drosophila* embryo^8^.

There are two major pathways of bulk mRNA degradation: 5’ to 3’ mRNA decay, mediated by the Xrn1 (Pacman (Pcm) in *Drosophila*) exonuclease, and 3’ to 5’ decay catalysed by the exosome.^9^ Phase separated cytoplasmic compartments called P-bodies contain many mRNAs and RNA-binding proteins, including Xrn1/Pcm, decapping factors and the conserved DEAD-box RNA Helicase DDX6 (Me31B in *Drosophila*).^10,11^ P-bodies are associated with mRNA degradation in yeast^12^, the early *Drosophila* embryo^7^ and neural crest cells in chick embryos.^13^ However, in other contexts, such as the *Drosophila* oocyte or mammalian cells, mRNAs are not degraded in P-bodies, but are instead stored and translationally repressed.^14,15^

In the early *Drosophila* embryo, the anterior-posterior (AP) axis is patterned by a hierarchical gene regulatory network, including gap, pair-rule and segment polarity genes.^16^ A highly studied gene in this cascade is the pair-rule gene *even-skipped* (*eve*), which encodes a transcriptional repressor required for segmentation. Transcriptional regulation of *eve* is well characterized, both in terms of the regulatory regions which dictate its 7-striped pattern^17,18^, and the transcriptional parameters that are modulated across the expression domain^3,19–21^. Here, we investigate the role of mRNA stability in shaping the *eve* expression pattern in early *Drosophila* development. By modeling *eve* transcription and mRNA data, we provide evidence that increased mRNA degradation at the edges of *eve* stripes 2 and 4 contributes to sculpting of the mature stripe patterns. We also demonstrate that altering *eve* mRNA numbers disrupts AP patterning in the *Drosophila* embryo. Together, these data highlight how spatial regulation of mRNA decay can refine developmental gene expression patterns.

## Results

### *eve* mRNAs are more colocalized with P-bodies at the edge of stripe 2

To study the contribution of mRNA stability to establishment of spatial gene expression patterns, we focused on the pair-rule gene *eve*. *eve* is initially transcribed in bursts by nuclei present in two broad domains. *eve* transcription then refines during nuclear cycle 14 (nc14), as nuclei in the interstripe regions stop transcribing *eve*, to produce the mature 7-stripe pattern.^22,23,21,19,3,20^ smFISH staining of *eve* in early to mid nc14 recapitulates this, as shown in the embryo schematics with nuclei coloured based on the presence of 1 or 2 *eve* transcription sites (TSs) (Fig. 1A, *eve* smFISH probe positions and underlying data are in Fig. S1A-B). For smFISH imaging, nc14 embryos were aged by measuring membrane ingression, indicating the extent of cellularization. For simplicity, embryos at different stages in mid nc14 are labelled with a single membrane ingression value (e.g. Fig. 1B, C), but ranges of ingression were defined for each time point of interest (see Methods). Focusing specifically on the refinement of stripe 2 in mid nc14 (8 μm membrane ingression), nuclei with active TSs are present at the anterior and posterior stripe borders (Fig. 1B). Despite containing an active TS, these nuclei outside of the mature stripe domain have low cytoplasmic mRNA numbers compared to the stripe center, suggesting that *eve* is rapidly degraded in these regions (Fig. 1B). Consistent with this, the distribution of total *eve* mRNAs is sharper than that of nuclear *eve* mRNAs (Fig. S1C). Later in nc14 (16 μm membrane ingression), *eve* TSs are no longer visible in the inter-stripe regions (Fig. 1C), in agreement with data from MS2 live imaging of *eve*, which demonstrated that nuclei in the interstripe regions burst until mid nc14.^19^

**Figure 1:**
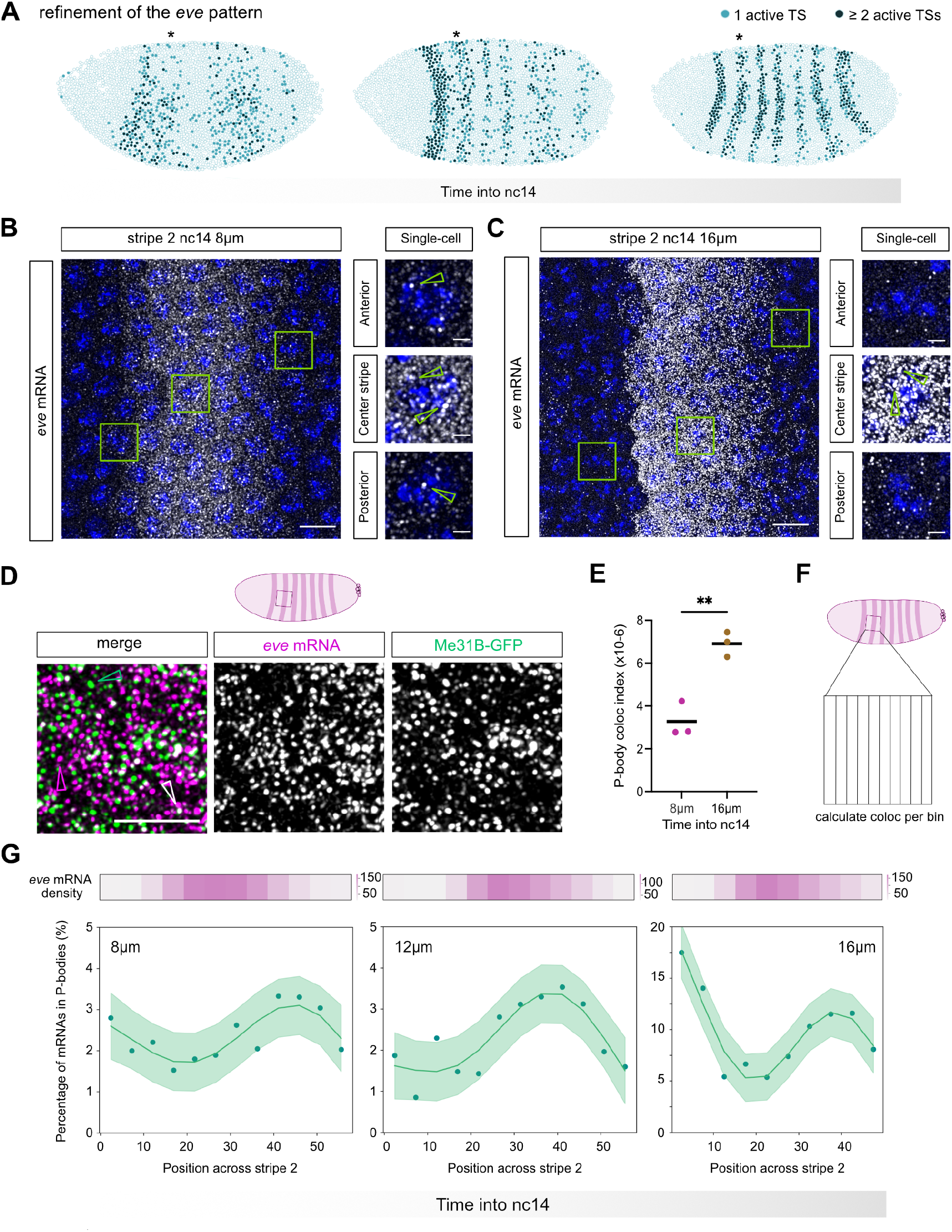
*eve* mRNAs are more colocalized with P-bodies at the edge of stripe 2. (A) Visualization of *eve* smFISH data from embryos at progressive time points during nc14, with nuclei coloured based on the number of active TSs (underlying data are in Fig. S1B). Asterisk indicates *eve* stripe 2. (B) Maximum projection of an smFISH image of *eve* stripe 2 in an embryo with 8 μm membrane ingression, with mRNAs shown in white and DAPI in blue. Green boxes mark cells displayed in the inset regions in the center and anterior and posterior edges. Active TSs were identified within nuclei in Z and are marked with a green arrow. (C) As in B, but embryo has 16 μm membrane ingression. (D) (i) Confocal image of a fixed nc14 Me31B-GFP embryo stained with smFISH probes for *eve* (magenta) and P-bodies (GFP, green). Images are projections of 7 slices; individual mRNAs (magenta arrowheads), P-bodies (green arrowheads), and colocalized mRNA and P-body signals (white arrowheads) are indicated. (E) Graph shows P-body colocalization index of *eve* mRNAs in mid nc14 embryos; *n* = 3 embryos, *p* = 0.0034, unpaired t-test used to determine significance with *α* = 0.05. (F) Schematic illustrating AP binning of a region across stripe 2 used in (G). (G) P-body colocalization data across stripe 2 for 3 timepoints from mid nc14. Data over the AP axis are fit with a gaussian process. The green shaded region marks the 95% confidence interval. Heatmaps above display the average mRNA per cell over the corresponding AP bins. Scale bars: 5 μm (main images), 1 μm (inset regions).

Based on the rapid refinement of the *eve* pattern, we hypothesized that spatially regulated mRNA stability could act in combination with transcriptional control to produce the mature *eve* pattern. As we previously showed that unstable mRNAs have a higher colocalization with P-bodies than stable mRNAs in the early embryo^7^, we used smFISH to calculate the colocalization of *eve* mRNAs with P-bodies. We focused on *eve* stripe 2 due to the in-depth study of its transcriptional regulation^18,22,24^. P-bodies were visualized using a fly stock carrying a Me31B-GFP protein trap^25^ (Fig. 1D). We calculated the P-body colocalization index, as described previously, which calculates the number of mRNAs colocalized with P-bodies while controlling for variation in mRNA and P-body numbers across embryos and developmental time.^7^ P-body colocalization was analysed across the whole stripe 2 domain, including the adjacent interstripe regions that show the characteristic sharp anterior stripe border and more graduated posterior border^26^, as in Fig. 1C. Comparing the P-body colocalization in 8 μm and 16 μm membrane ingression embryos demonstrates that *eve* mRNAs become more localized to P-bodies as the stripe refines (Fig. 1E), potentially suggesting a role for mRNA decay in stripe refinement.

The localization of *eve* mRNAs in P-bodies was then calculated in AP bins of ∼1 cell width across the stripe 2 domain (Fig. 1F). As P-bodies are mostly uniform across stripe 2, we calculated the percentage of colocalized mRNAs across the stripe. This analysis revealed that mRNAs in the interstripe regions are more colocalized with P-bodies than mRNAs in the center of stripe 2 (Fig. 1G, Fig. S1D-E). The strength of this effect increases as nc14 progresses (Fig. 1G), consistent with the overall increase in P-body colocalization over time (Fig. 1E). These data suggest that *eve* mRNAs in the interstripe regions may be less stable than those in the stripe center. Although we previously used separate 5’ and 3’ smFISH probe sets to detect potential mRNA degradation intermediates^7^, this approach was not feasible for the short *eve* mRNA. Therefore, as the determinants of P-body targeting remain incompletely understood, increased colocalization may also reflect other properties, such as transcript age, i.e. time since synthesis (see below).

### Modeling mRNA degradation reveals different decay kinetics across *eve* stripe 2

We next studied how mRNA degradation contributes to the refinement of the *eve* stripe 2 domain by comparing mRNA numbers with transcriptional activity. Previously published MS2-MCP live imaging data showed that some *eve* stripes, e.g. stripes 2 and 4, are transcriptionally refined by 20 mins into nc14 (Fig. 2A), and the full 7-stripe pattern is refined by 25 mins into nc14.^19^ We therefore focused on the 20-minute time point, up to which the refinement of stripe 2 is occurring. Nuclei spanning the region between the posterior edge of stripe 1 and the anterior edge of stripe 3 were selected in the live dataset, and the summed MS2 fluorescence up to this time point was calculated (transcription output, Fig. 2B). *eve* mRNA numbers in the corresponding region were quantitated using *eve* smFISH staining in fixed embryos (total mRNA, Fig. 2B, S2B), which were carefully staged in nc14 using membrane ingression.^23,27,28^ Live and smFISH data across stripe 2 (boxed region in Fig. 2B) were binned into a 5 x 5 grid across the AP and DV axes, and data within each bin were averaged (see Methods). Comparison of the resulting heatmaps demonstrates that the mRNA data is more refined than the summed transcriptional output, particularly at the anterior border where the transcriptional output is higher than in the posterior (Fig. 2B). These data further suggest that post-transcriptional regulation contributes to refinement of the *eve* stripe 2 expression pattern.

**Figure 2:**
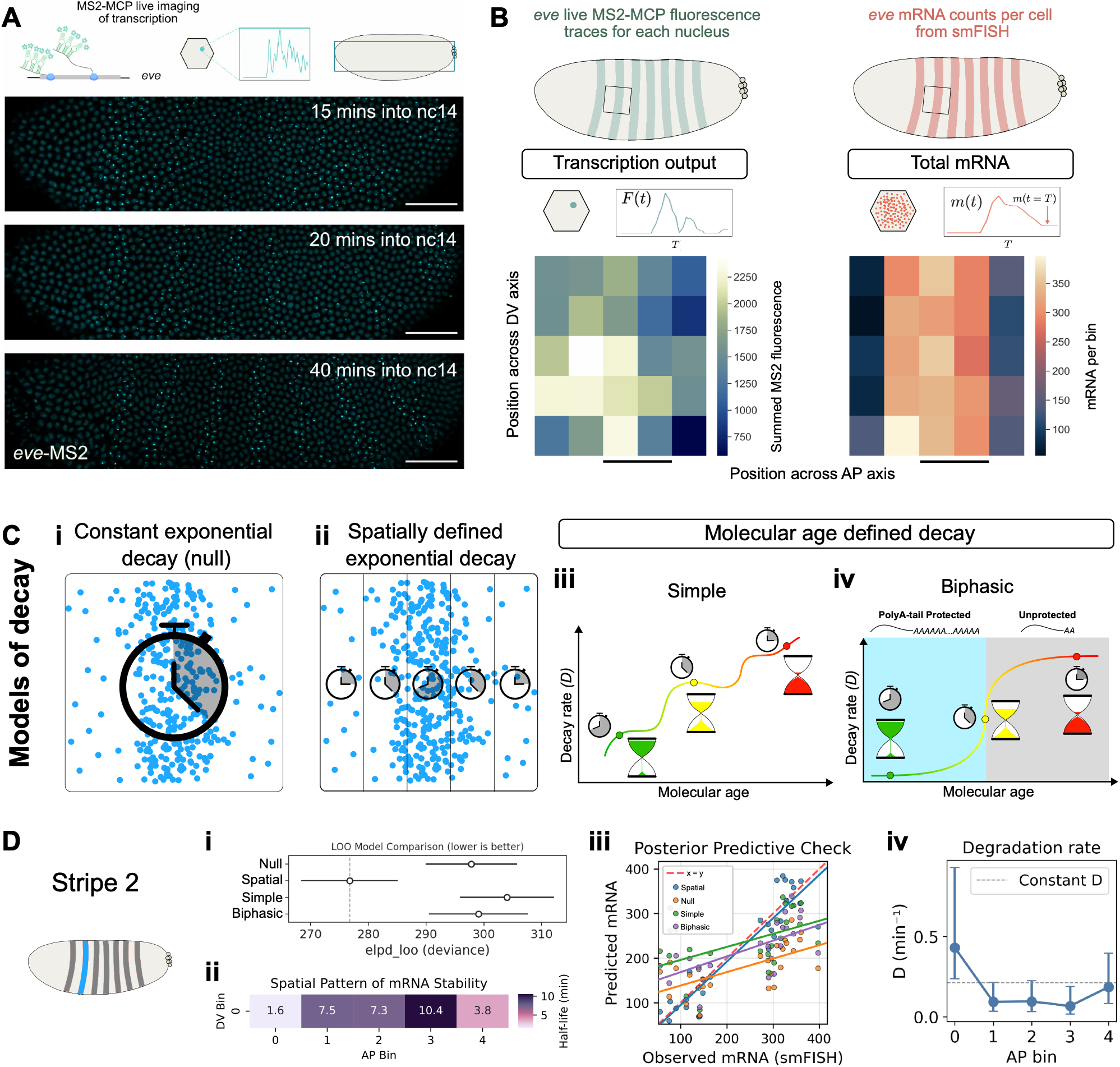
Modeling mRNA degradation reveals different decay kinetics across *eve* stripe 2. (A) Schematic of MS2-MCP-GFP detection of *eve* transcription in each nucleus. Stills from a live movie from ^19^ at 15, 20 and 40 mins into nc14 showing refinement of the *eve* pattern. (B) Embryos marked with regions for comparison of *eve* live imaging MS2 data (green) and smFISH data (orange). Schematics depict the MS2 trace and the total mRNA across time. Heatmaps showing the average summed transcriptional output (left) and mRNA per bin (right) in AP and DV bins up to 20 mins into nc14 for stripe 2. The solid line marks the position of the mature stripe. (C) Schematics depicting: i) the null model where decay is constant across space and time, ii) spatially defined decay along the AP axis, iii) the simple molecular-age model where the decay can vary with age as modelled by a gaussian random walk, and iv) the biphasic molecular-age model inspired by polyA-tail protection. (D) Results from the four models for stripe 2 (coloured blue). (i) Comparison of the models’ performance on unseen data by performing LOO cross validation. The lower the expected log predictive density (epld_loo), the better. Bars represent the standard error of the ELPD estimate. (ii) Heatmap showing the median half-life value for each of the AP bins from the spatial model. (iii) Scatter plot comparing the observed mRNA to the predicted mRNA when the median of the learned posterior distribution for each parameter is used in the model to process the MS2 traces. (iv) Graph showing the median and 95% confidence intervals (CI) for the estimated decay rate for each of the five AP bins from the spatial model. The dotted horizontal line represents a constant decay rate across space. The line is placed at the midpoint between the maximum of the lowest 95% CI values and the minimum of the highest 95% CI values across the five AP bins.

Theoretically, a few different methods of decay could produce the *eve* stripe 2 pattern. We built four simple mathematical models to infer degradation rate and then compared the performance using leave-one-out (LOO) cross validation, to identify which method of decay best fits the data. Conceptually, the four models are as follows: (1) The null model is a simple exponential decay model that assumes decay is constant across molecular age and space (Fig. 2Ci). (2) A spatially dependent model where decay varies along the AP axis but is not influenced by transcript age or the DV axis (Fig. 2Cii). (3) A simple transcript age dependent model where decay generally gets stronger as the age of the molecule increases, but there is no spatial influence (Fig. 2Ciii). (4) A biphasic transcript age dependent model that assumes no spatial influence but attempts to mimic the slow decay phase of a protective poly-A tail followed by a fast decay phase after a poly-A tail has been shortened^29,30^ (Fig. 2Civ). Mathematically, the evolution of mRNA numbers over time for all four models is described by the ordinary differential equation (ODE):

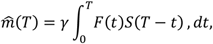

where cytoplasmic mRNA numbers 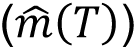 are established by a balance of transcription (*F*(*t*)) and mRNA decay (*S*(*T* − *t*)). In this model, the transcription dynamics are described by the fluorescent MS2 signal over time (*F*(*t*)), scaled by a conversion factor (*γ*) which converts the fluorescent signal into the number of mRNA molecules produced. *γ* is inferred in the model and is shared across all bins in the expression domain. This equation describes the mRNA at time *T*, depicted as *m*(*t* = *T*), which corresponds to the mRNA counts from the smFISH data. The only place these four models differ, is in their definition of the decay kernel *S*(*T* − *t*). See Methods for full details.

These models are constructed around several assumptions which are outlined as follows. Firstly, the process of transcriptional regulation is assumed to follow a first-order process represented by the ODE above, as is standard for models of transcriptional regulation.^7,31^ Secondly, the conversion factor between fluorescence and mRNA number is assumed to be constant across all regions in the expression domain, which we believe to be a fair assumption and is typically assumed for this type of data.^3,4^ Thirdly, the stripe’s peak transcription in the MS2 trace is at the same position as the peak in the total mRNA as captured by smFISH to facilitate data alignment. Finally, and uniquely to the spatially dependent model, we assume that regions (bins) that share a given AP position also share the same degradation rate. That is, we allow for variation in mRNA stability across the AP axis but assume that it is constant across the DV axis. This is a specific assumption appropriate for the expression pattern of *eve* which is dynamic across the AP axis and mostly invariant in DV, typical for an AP patterning factor (Fig. S2A).

The models were fit to the real *eve* mRNA data from Fig. 2B using Markov chain Monte Carlo (MCMC) methods. Results indicate that the spatial decay model outperforms the null and two molecular age models in the LOO comparison (Fig. 2Di). This model demonstrates that mRNA decay is modulated across *eve* stripe 2, with an inferred half-life of >7 mins at the stripe center and <4 mins in the interstripe regions (Fig. 2Dii). We performed a posterior predictive check to determine how well each model can recapitulate the experimentally measured mRNA using the learned parameters for each model. This reveals that only the spatial model can produce mRNA values that match the full range observed in the experimental data (Fig. 2Diii). We then visualized the 95% confidence intervals for the predicted decay rates for each AP bin, which shows that a constant decay cannot satisfy the intervals for all five bins (Fig. 2Div). The average *eve* half-life across all five bins is 6.1 min, which corresponds closely to a previous estimate of 6.5 min for *eve* mRNAs, based on measurements of mRNA levels by Northern blot following transcriptional inhibition.^32^

To control for potential temporal misalignment of the smFISH image, a slightly younger embryo (7 μm membrane ingression) was used (Fig. S2B). It is possible that this embryo is a better temporal match to the MS2 data, which were generated from embryos carrying a BAC with *MS2-yellow* in place of the *eve* coding sequence, adding ∼5.5kb to the transcribed sequences.^19^ Analysis of this temporally shifted embryo shows similar spatial modulation of *eve* stability across stripe 2. Additionally, to control for potential spatial misalignment, greater than that corrected by our binning approach, the MS2 data were shifted half a bin width to the left and right and then reanalysed, again showing consistent results with our initial alignment (Fig S2B). Together, these data support spatial regulation of *eve* stability across stripe 2, with ∼2-5-fold difference in the half-life of *eve* between the interstripe regions and the center of stripe 2.

We next tested the ability of the model to recover known parameters using the MCMC method to ensure the accuracy of the learned decay rates. We selected different decay values for each of the five AP bins and combined them with the average transcription trace for each bin to generate *in silico* mRNA data (Fig. S3A). We then trained the spatial model using this *in silico* data to determine if the model could capture the predefined decay values. The results of this are shown in Fig. S3B-C and demonstrate that inferred parameters correspond well to the true values.

### *eve* stripe 3 may not require spatial decay like stripes 2 and 4

We next examined whether spatially regulated decay also contributes to the generation of *eve* stripes 3 and 4. Using smFISH measurements of total mRNA abundance (Fig. S4) together with MS2 transcriptional data, we fitted the same set of MCMC models. Interestingly the results for stripe 3 suggest that all four models can fit the data equally well (Fig. 3A). The LOO comparison shows similar deviance for all four models with large overlap in the standard error bars of the ELPD estimate (Fig. 3Ai). The inferred half-lives for the spatial model are shown in Fig. 3Aii. Posterior predictive checks revealed that each model accurately reproduced the experimental mRNA numbers (Fig 3Aiii). A constant decay rate easily overlaps the 95% confidence intervals of the decay rates for each AP bin predicted by the spatial model (Fig. 3Aiv). This suggests that, although a spatial model is possible, it is not required to explain the experimental data for stripe 3. In contrast, stripe 4 resembled stripe 2, with the spatial decay model providing a significantly better fit to the experimental data (Fig. 3Bi–iv). The inferred half-life is >7 mins at the center of stripe 4 and <2 mins in the interstripe regions (Fig. 3Bii).

**Figure 3:**
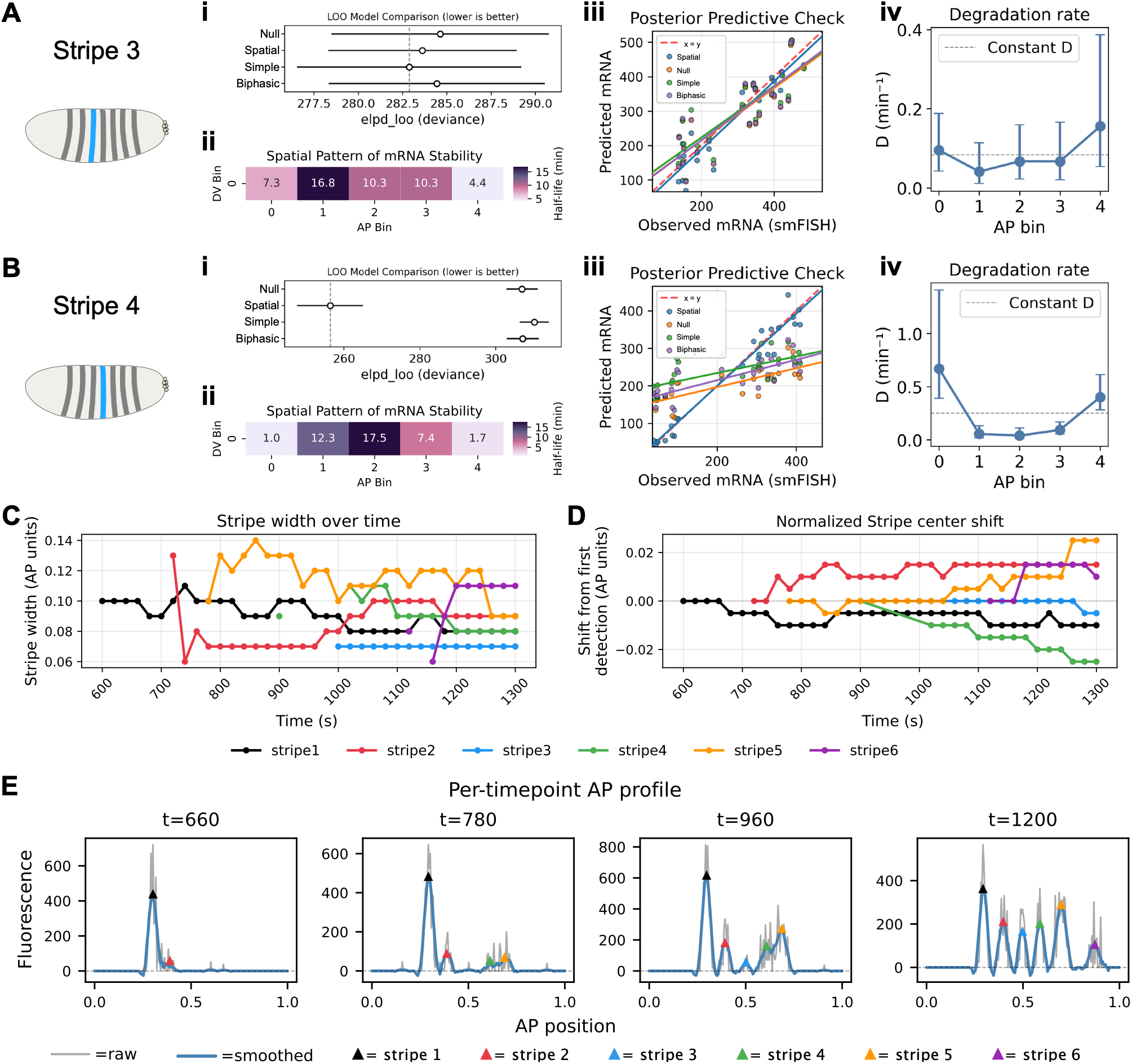
*eve* stripe 3 may not require spatial decay like stripes 2 and 4. (A-B) As in Fig. 2D, but the data are for *eve* stripes 3 (A) and 4 (B). (C) Line plot of the stripe widths over time, the colour key for each stripe is shown below the graph. (D) Line plots depicting stripe center position (shift) across time, normalized to their first detected position. Stripe centers and widths were programmatically identified from the MS2 data as described in the Methods. (E) Graphs show the MS2 fluorescence, as raw and smoothened traces, at specific time points (in seconds) in nc14 that highlight how stripes 2 (red), 3 (blue), and 4 (green) emerge.

To understand why stripe 3 behaves differently from the other two stripes tested, we studied the stripe morphology over time starting with stripe width. At our measured time point (1200s), stripe 3 is thinner than the other visible stripes and has not changed in width relative to earlier times (Fig. 3C). We then visualized the stripe position over time to determine how much each stripe shifts along the AP axis during stripe refinement. Once again, at the 1200s time point, stripe 3 has not shifted position whereas all other visible stripes have shifted from their first point of detection (Fig. 3D). This likely reflects the different way that stripe 3 emerges, which is in an area between the 2 initial broad domains of eve expression, rather than progressively refining from the broad pre-pattern as is the case for stripes 2 and 4 (Fig. 3E, 1A). Taken together, this suggests that stripes 2 and 4 shift position across time and are wider in general, leading to residual *eve* transcripts at the edges of the stripes that need to be cleared to define a sharp stripe boundary. In contrast, stripe 3 does not shift position in our time window and stays narrow avoiding residual expression at the edges that needs to be removed.

As described for stripe 2, we performed temporal and spatial shifts for *eve* stripes 3 and 4. In all cases the spatial model outperforms the others for stripe 4 (Fig. S4D), indicating it is robust to spatial and temporal perturbations, similar to stripe 2 (Fig. S2B). However, while the temporal shift for stripe 3 shows that the null model is sufficient to explain the data, the spatial model better fits the data following the spatial shifts (Fig. S4C). As described above, stripe 3 is narrower, less mobile, and does not refine away from other stripes to the same extent as stripes 2 and 4 (Fig. 3E). Based on this, we speculate that the spatial shifts realign the MS2 transcription and mRNA smFISH profiles in such a way that it more resembles the conditions of stripes 2 and 4. Thus, it is predictable that spatially varying decay would be needed to generate the observed mRNA numbers. Therefore, while it remains possible, we favour the conclusion that spatially varying decay is not needed for stripe 3. In contrast, stripes 2 and 4 are robust to the spatial perturbations, which suggests spatially varying decay is necessary for these stripes’ refinement rather than being a consequence of misalignment.

### *In silico* manipulations with uniform *eve* mRNA stability disrupts stripe 2 and 4 patterns

To better understand the requirement for spatial variation in *eve* mRNA decay, we investigated the effect of uniform *eve* mRNA stability on stripe refinement *in silico*. Using the spatial models trained on stripes 2, 3, and 4, we generated synthetic mRNA data from the *eve* transcription traces, analogous to the posterior predictive checks (Fig. 2Diii and Fig. 3Aiii, Biii), but with a uniform half-life value for all AP bins. We first used the model to synthesize mRNA values using a short half-life value for all AP bins calculated by averaging the half-lives from the left and right edge bins from the fitted spatial model. We then repeated this using the longest half-life inferred for each stripe from the fitted spatial model. The results for both the short and long half-lives were then compared to the real mRNA data observed for stripes 2-4 at the 20 min timepoint (Fig. 4A-Ci). For *eve* stripes 2 and 4, with the long half-lives (10 min for st 2 and 18 min for st 4), the center of the stripe forms a peak which is similar to the observed data (Fig. 4Aii-iii, 4Cii-iii). However, at the stripe edges, higher mRNA numbers are observed than in the real data, resulting in a broader stripe (Fig. 4Aii-iii, 4Cii-iii). For the short half-lives (3 min for st 2 and 1 min for st 4), the interstripe regions are cleared of mRNA. However, the center of the stripe fails to form the strong peak in *eve* mRNA numbers observed in the real data (Fig. 4Aii-iii, 4Cii-iii), again resulting in a failure to clearly define a stripe. In contrast, the mRNA data from the spatial model matches the full experimental data (Fig. 4Aiii, Ciii), as described above (Fig. 2Diii, 3Biii).

**Figure 4:**
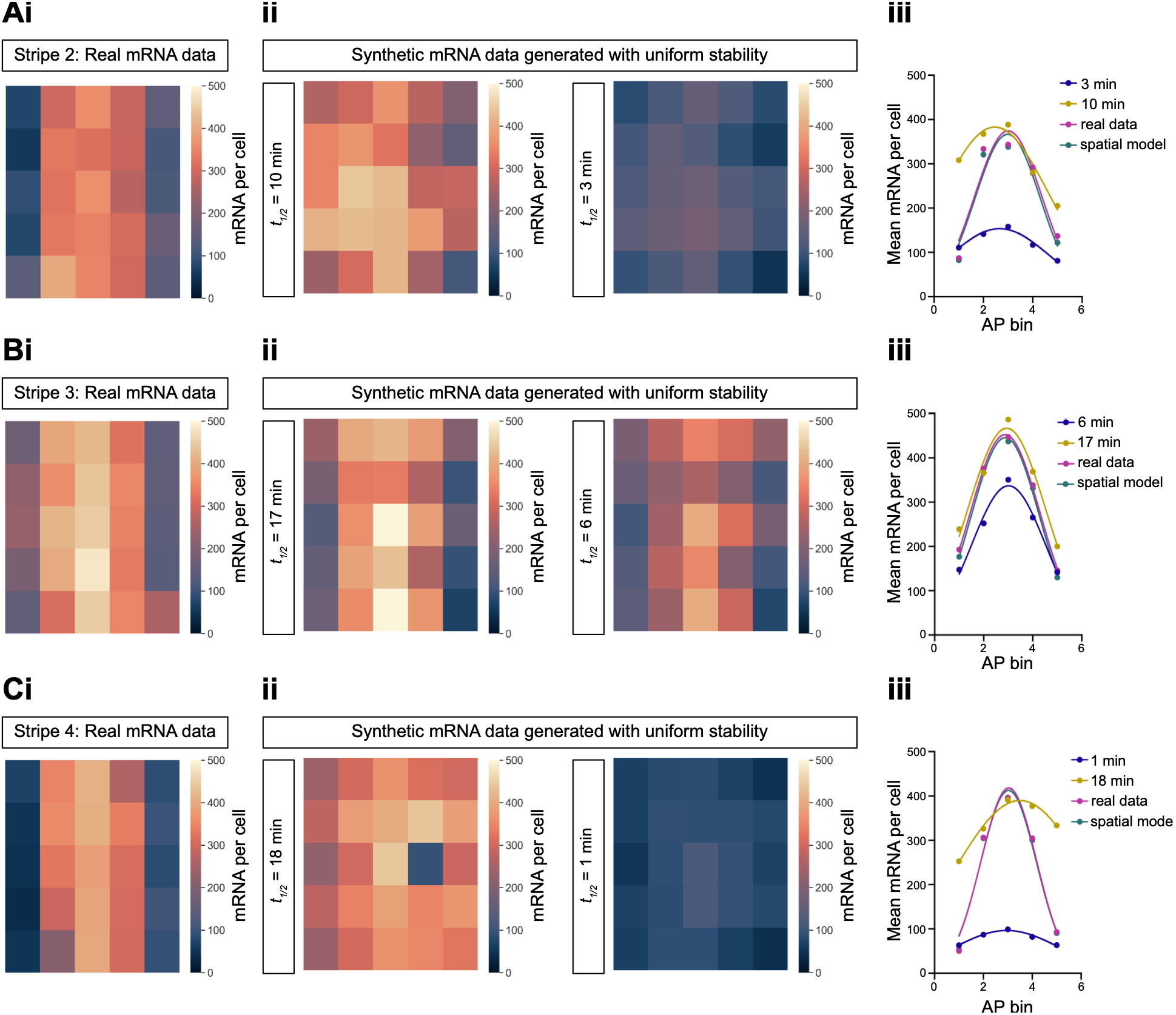
*In silico* manipulations with uniform *eve* mRNA stability disrupts stripe 2 and 4 patterns. (A-C) Synthetic mRNA data generated from modeling *eve* transcription data up to 20 minutes in nc14 with constant half-life across stripe 2 (A), stripe 3 (B) and stripe 4 (C). (Ai-Ci) Heatmaps depict the average summed mRNAs in AP and DV bins from *eve* smFISH images (also shown in Fig. 2B, S4A, B, but rescaled here for cross comparison). (Aii-Cii) Heatmaps showing the synthetic mRNA counts generated with the longer and short half-lives for *eve*, as indicated to the left of the heatmap. (Aiii-Ciii) Mean mRNA numbers per cell summed across all DV positions in each AP bin, for the real and simulated data, fit with a gaussian distribution.

Analysis of *eve* stripe 3 reveals that a stripe is defined either with uniform shorter (6 min) or longer (17 min) half-lives, although the shorter half-life reduces mRNA numbers in the stripe compared to the real mRNA data that is particularly evident in the stripe center (Fig. 4Bi-iii). The mRNA data from the spatial model (Fig. 3Aiii) are also shown as a comparison (Fig. 4Biii). These results are consistent with uniform mRNA stability being sufficient to define *eve* stripe 3, as described and explained above (Fig. 3). Overall, these simulations show that spatially regulated mRNA decay across stripes 2 and 4 is required to refine the mature stripe pattern and produce sharp boundaries.

### Swapping the *eve* 3’UTR alters *eve* mRNA numbers and formation of ectopic stripes

The analysis above (Fig. 4) shows how uniform *eve* mRNA stability alters the number of *eve* mRNAs and disrupts definition of stripes 2 and 4. In the absence of a known regulator of *eve* mRNA degradation, it is currently technically unfeasible to manipulate *eve* mRNA stability spatially *in vivo*. Therefore, to extend our findings *in vivo*, we instead investigated whether *eve* mRNA numbers are important for AP patterning. To this end, we aimed to alter *eve* mRNA stability by changing its UTR, as sequences in the 3’UTR play a major role in regulating mRNA stability.^33^ A previous study has shown that in a transgenic reporter with the *eve* stripe 2 enhancer and promoter, the presence of the *α1-tubulin* 3’UTR alters the temporal profile of mRNA accumulation compared to that observed with the *eve* 3’UTR. Specifically, the reporter with the *α1-tubulin* 3’UTR results in fewer mRNAs early in nc14, then comparable levels, followed by increased mRNA accumulation in late nc14 embryos.^34^ Based on this finding, we hypothesized that swapping the *eve* 3’UTR for the *α1-tubulin* 3’UTR would alter the number of *eve* mRNAs in the embryo and allow us to test the effect on AP patterning. Therefore, we used CRISPR-Cas9 scarless genome editing to replace the endogenous *eve* 3’UTR (179 bp) with that of *α1-tubulin* (256 bp) (Fig. 5A). The resulting *eve-tubulin* flies were found to be homozygous lethal.

**Figure 5.**
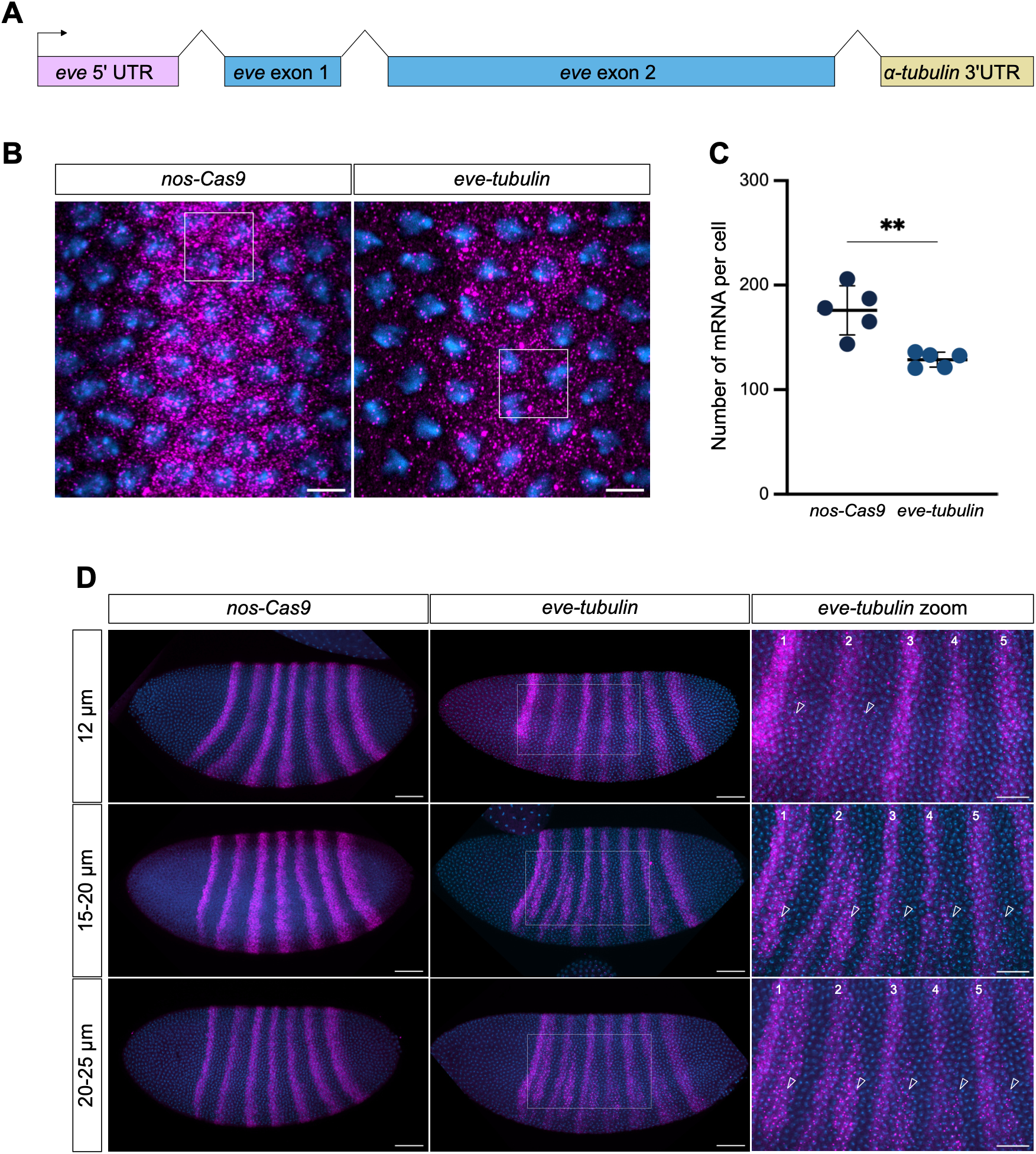
Swapping the *eve* 3’UTR results in altered *eve* mRNA numbers and formation of ectopic stripes. (A) Schematic showing the *eve* gene with the *α1-tubulin* 3’UTR. (B) Maximum projections of smFISH images of *eve* stripe 2 in *nos-Cas9* and *eve-tubulin* nc14 embryos at 8 μm membrane ingression. *eve* mRNAs are shown in magenta, nuclei (DAPI) in blue. The white boxes mark the area used for analysis. Scale bar: 5 μm. (C) Graph shows the number of *eve* mRNAs per cell in *nos-cas9* and *eve-tubulin* embryos. n=5, mean±s.d., \*\**P<*0.01 (unpaired two-tailed *t*-test). (D) Maximum projections of *eve* smFISH staining of *nos-Cas9* and *eve-tubulin* in progressively older nc14 embryos. Higher magnification zooms of the boxed regions are shown. *eve* stripes are numbered, arrowheads indicate the ectopic stripes. Scale bars: 50 μm whole embryos, 25 μm, zoom.

To understand this lethality, we first used smFISH to image *eve* stripe 2 in *eve-tubulin* and control *nos-Cas9* mid nc14 embryos. In all the embryo stainings presented in this study, *lacZ* smFISH probes were also included, so that homozygous *eve-tubulin* embryos (hereafter referred to as *eve-tubulin* embryos) could be identified based on the absence of a balancer chromosome that carried *ftz-lacZ*. Quantitation of *eve* mRNA numbers in the center of stripe 2 reveals a significant reduction in the *eve-tubulin* nc14 embryos (∼8 um membrane ingression), with ∼30% less *eve* mRNAs per cell (Fig. 5B, 5C). This suggests that swapping the UTR to the *α1-tubulin* 3’UTR reduces the stability of the *eve* mRNAs. To assess protein levels, we combined *eve* smFISH with Eve antibody staining (Fig. S5A). As the Eve antibody staining was noisy, we quantified the fluorescent mRNA and protein signals across all seven *eve* stripes from whole embryo images (Fig. S5A). This semi-quantitative analysis suggests that, overall, there is an average reduction of ∼35% in both mRNA and protein in the *eve-tubulin* embryos compared to the control (Fig. S5B-C). Together, these data support a reduction in *eve* mRNA numbers and protein levels in mid-nc14 *eve-tubulin* embryos, as suggested by the findings from the different UTR-containing *eve* reporters.^33^

Visualization of the *eve* stripes in embryos using smFISH at different stages of nc14 reveals that ectopic secondary *eve* stripes emerge in the *eve-tubulin* embryos (Fig 5D). Transcription of ectopic *eve* stripes is initially detected between stripes 1-2 and 2-3 at 12 μm membrane ingression, then additional ectopic stripes become more obvious throughout the *eve-tubulin* embryos at later stages in nc14 (Fig. 5D). As described for the secondary *eve* stripes in wildtype embryos, which are not detectable in embryos until the onset of gastrulation^35,36^, the ectopic stripes are initially more evident ventrally (Fig. 5D). As well as the earlier activation in the *eve-tubulin* embryos, the secondary *eve* stripes are stronger and positioned closer to the preceding major *eve* stripe in the interstripe region (Fig. 5D). In contrast, the minor *eve* stripes in wildtype embryos have very weak expression, typically undetectable by fluorescent in situ hybridization^37^, and are centered in the interstripe region.^35,36^ Eve antibody staining shows accumulation of Eve protein in the ectopic secondary stripes in *eve-tubulin* nc14 embryos (Fig. S5D).

### Disrupted AP gene regulation and patterning in *eve-tubulin* embryos

To study how lower *eve* mRNA numbers leads to ectopic *eve* stripes, we used smFISH to detect *eve* and other pair-rule mRNAs, including *odd-skipped* (*odd*), *runt* and *sloppy paired* (*slp*). Eve represses *odd* transcription^37–39^ (Fig. 6A), and double *eve* and *odd* smFISH stainings in nc14 embryos reveals that the *odd* stripes are expanded in the *eve-tubulin* nc14 embryos (Fig. 6Bi, ii). Odd represses *runt* transcription^37,39,40^ (Fig. 6A), and the *runt* stripes are thinner in *eve-tubulin* nc14 embryos compared to the control (Fig. 6Ci, ii). These data are consistent with the weaker Eve stripes leading to an expansion of Odd, which results in lower Runt levels. As Runt represses *eve*^37,39,41^, we suggest that the reduced Runt levels allow earlier transcription of the secondary stripes. In addition, the double *runt eve* staining shows that *runt* stripes abut the anterior border of the secondary *eve* stripes (Fig. 6Cii). This suggests that the thin Runt stripes establish the anterior border of the ectopic *eve* stripes, consistent with Runt’s known role as a repressor of *eve* transcription.^37,39,41^ Eve also represses *slp* transcription^37,39,42^ (Fig. 6A), and double *eve* and *slp* smFISH stainings reveals that the *slp* stripes are broader in the *eve-tubulin* nc14 embryos (Fig. 6Di, ii), consistent with less Eve protein compared to the control embryos (Fig. 5). As the *slp* stripes abut the posterior of the ectopic *eve* stripes (Fig. 6Dii), and Slp represses *eve* transcription^37,39,42^, Slp appears to define the posterior border of the ectopic *eve* stripes.

**Figure 6.**
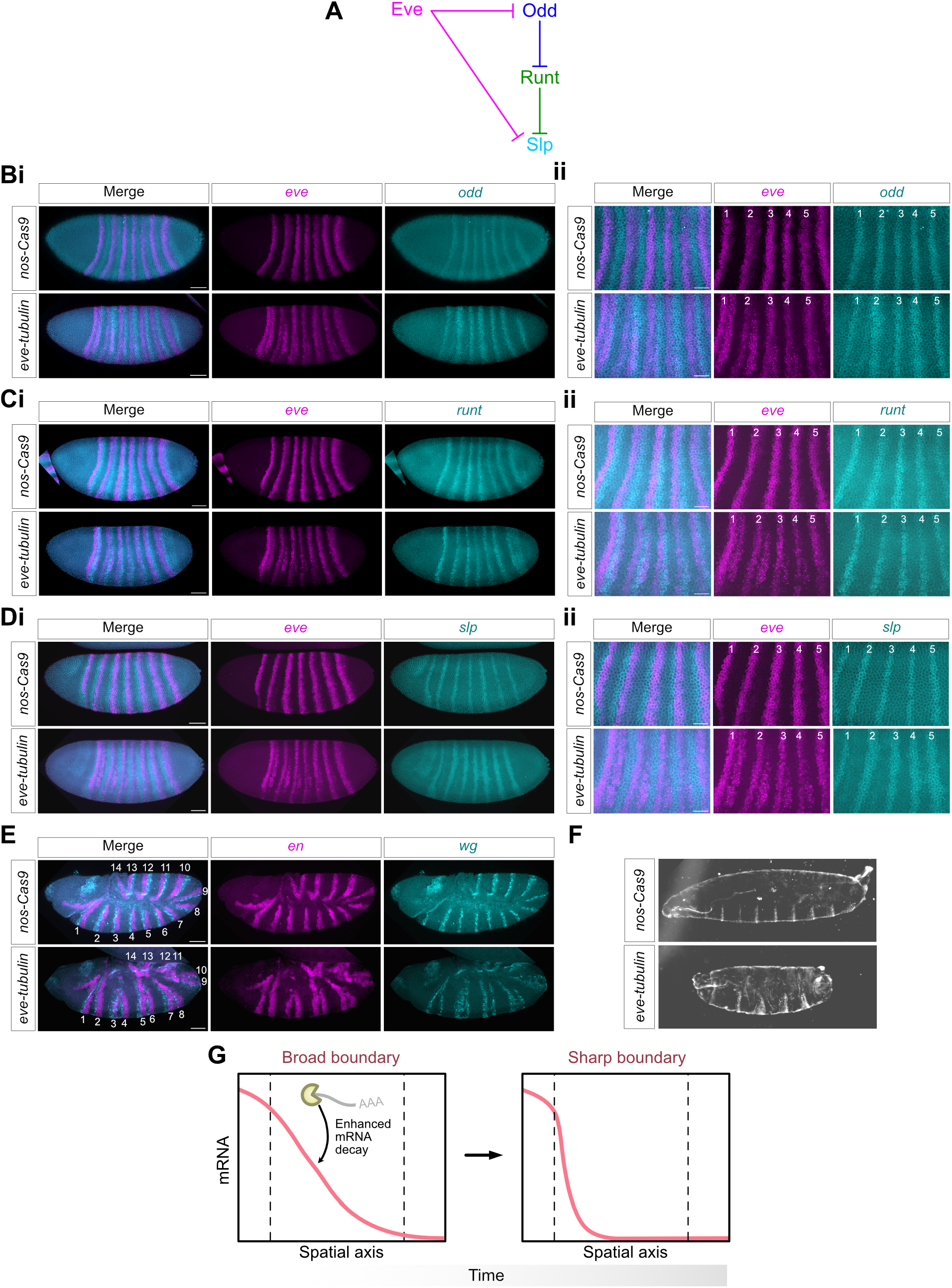
*eve-tubulin* embryos have disrupted AP gene regulation and a pair-rule phenotype. (A) Schematic showing repressive interactions (blunt arrows) between Eve, Odd, Runt and Slp, figure simplified from ^39^. (B) Maximum projection smFISH images of *nos-Cas9* and *eve-tubulin* nc14 embryos (17-20 μm membrane ingression) stained with *eve* and *odd* probes. Whole embryo (Bi) and higher magnification views with the stripes numbered (Bii), shown as merged images (left panel) and single channel images (middle and right panels). (C-D) As in (B) except that the embryos are stained with *eve* and *runt* (Ci, ii) or *eve* and *slp* probes (Di, ii). Scale bars: 50 μm whole embryos, 25 μm, zoom. (E) Maximum projections of smFISH staining of stage 9 *nos-Cas9* and *eve-tubulin* embryos with *en* and *wg* probes. The 14 stripes are numbered. (F) Images of *nos-Cas9* and *eve-tubulin* cuticles. (G) Schematic demonstrating how enhanced mRNA decay across a broad transcriptional boundary can result in a sharp boundary at a later timepoint.

Given the altered expression patterns of *eve*, *odd*, *runt* and *slp* in the *eve-tubulin* nc14 embryos, we next visualized the downstream segment polarity genes, *engrailed* (*en*) and *wingless* (*wg*). In wildtype embryos, the anterior border of *fushi tarazu* (*ftz*) is one cell broader than that of *odd*, due to Runt and Odd-paired (Opa) repression of *odd*. This offset, which allows the even-numbered *en* stripes to be transcribed^37^, is less evident in *eve-tubulin* nc14 embryos (Fig. S6A), and the *en* and *wg* stripes are disrupted (Fig. 6E). Analysis of the *en* stripe positions relative to the cephalic furrow reveals that it is an anterior shift of the even numbered *en* stripes (Fig. S6B), the ones that overlap with the *eve* minor stripes in a wildtype embryo.^36,37^ Consistent with this shift in *en* and *wg* expression, analysis of the cuticles from *eve-tubulin* embryos shows a pair-rule phenotype (Fig. 6F). This phenotype is characteristic of *eve* hypomorphs, whereas the *eve* null mutants lack segmentation within the trunk of the embryo.^43,44^ We observe the pair-rule phenotype in 25-30% of the embryos, similar to the proportion (25%) of expected homozygous *eve-tubulin* embryos. Therefore, the disrupted segmental patterning can explain why *eve-tubulin* homozygotes are inviable. Together, the results presented here suggest that the *eve* mRNA numbers/cell, which are established by spatial regulation of mRNA stability, are critical for developmental patterning. The altered *eve* mRNA accumulation profiles in nc14 *eve-tubulin* embryos perturbs repressive interactions with other pair-rule genes, resulting in misregulation of downstream gene expression (e.g. *en*) and embryonic segmentation.

## Discussion

In this study, our mathematical modeling of live *eve* transcription and smFISH data provides evidence that, like transcription parameters, mRNA degradation rates can vary between cells across expression domains. *eve* mRNA stability shows spatial variation that helps refine the stripe patterns in early to mid nc14 embryos. The model-inferred mean decay rate is ∼8 mins for *eve* stripes 2-4, which agrees well with a previous estimate of ∼6.5 minutes^32^, with a ∼2-5 fold modulation of *eve* mRNA half-life across stripes 2 and 4. We observe higher colocalization of *eve* mRNAs and P-bodies at the edges of stripe 2 than the center. As we previously provided evidence that 5’ to 3’ mRNA degradation occurs in P-bodies in the early *Drosophila* embryo^7^, one interpretation is that the altered *eve* mRNA colocalization in P-bodies across the stripe reflects spatial differences in mRNA stability.

Changes in P-body localization may also reflect factors other than mRNA stability. We hypothesized that older mRNAs might preferentially associate with P-bodies, for example through increased mRNA damage or modification. Although mRNA age is difficult to assess *in vivo*, there are precedents for aberrant or modified mRNAs being localized to P-bodies. mRNAs with a premature stop codon, which are subject to nonsense-mediated decay, are targeted to P-bodies in yeast.^45^. Targets of CDS-m6A decay, triggered by m6A modification in the coding sequence, are also enriched in P-bodies.^46^ Time-varying decay is conceptually attractive as it would enable self-refinement of transcriptional pre-patterns, since transcripts from cells that have ceased transcription would be degraded more rapidly. However, for *eve* stripes 2 and 4, spatially varying decay provided a better fit to the experimental data than the two age-dependent models tested.

Although we considered spatial and temporal variation in *eve* mRNA degradation separately, the dynamic shifts in *eve* expression during early embryogenesis mean that spatial changes in *eve* mRNA decay will likely also lead to temporal changes. Previous analyses of *eve* stripe 2 using MS2 and smFISH data in late nc14 assumed and validated a constant mRNA decay rate across the stripe.^3^ We propose that by focusing on stripe refinement we detect spatial differences in stability, whereas such regulation is not evident later in nc14 once interstripe transcription ceases and the mature pattern is established. Together, these observations support both spatial and temporal regulation of *eve* mRNA stability during nc14. Temporal regulation of mRNA decay has been suggested for *ftz*, as the mRNA half-life decreases from ∼14 minutes in nc13 to shorter values that remain relatively constant (8, 7 and 6 min) across nc14 time points.^47^ More broadly, developmental changes in mRNA stability, including longer half-lives at later stages, have been reported.^5,48^ It will be interesting to test spatiotemporal regulation of *eve* mRNA stability, and more generally of other patterning mRNAs, in future experiments (see Limitations).

A key unresolved question is how differential mRNA stability is generated across *eve* stripes. One possibility is regulation by RNA-binding proteins, which typically bind consensus motifs in 3’UTRs.^33^ mRNA capture studies in early *Drosophila* embryos have identified numerous novel RNA-binding proteins, suggesting that there is extensive uncharacterized post-transcriptional regulation.^49,50^ miRNAs are another non-mutually exclusive option for regulation of mRNA stability. Many microRNAs display spatially restricted gene expression patterns in the *Drosophila* embryo^51^, and modeling suggests that interactions between miRNAs and targets in adjacent domains can sharpen expression boundaries.^52,53^

Using *in silico* simulations, we showed that uniform *eve* mRNA stability disrupted the *eve* stripes. Although spatial manipulation of *eve* mRNA stability *in vivo* is not currently feasible, previous studies with a reporter containing the *eve* enhancer and promoter demonstrated that replacing the *eve* 3’UTR with the *α1-tubulin* 3′UTR altered mRNA accumulation dynamics.^34^ Consistent with this, swapping the *α1-tubulin* 3’UTR into the endogenous *eve* locus reduced *eve* mRNA and protein levels by ∼30% in early nc14 embryos. These *eve-tubulin* embryos displayed strong, ectopic activation of minor *eve* stripes at asymmetric interstripe positions, leading to shifts in pair-rule (*odd, slp,* and *runt*) and segment polarity (*en* and *wg*) gene expression and a pair-rule phenotype. Therefore, while loss of the minor *eve* stripes does not disrupt parasegment organization^42,54^, our data show that premature activation of these stripes disrupts AP patterning.

The strong, early activation of the minor *eve* stripes in *eve-tubulin* mid nc14 embryos was unexpected, as these stripes are only weakly and transiently expressed in wildtype gastrulating embryos.^35,36^ Previous reporter studies identified a 1.7 kb fragment, starting 37bp downstream of the *eve* 3’UTR, that drives equivalent expression to the minor stripes in embryos undergoing germ band elongation. However, this activity is suppressed when combined with an upstream regulatory fragment containing late primary *eve* stripe enhancer activity, suggesting that correct minor stripe expression depends on interactions between 5’ and 3’ regulatory sequences.^55^ It is possible that our UTR swap disrupted this interaction, leading to the strong, early minor *eve* stripe expression we observed. We consider this unlikely for 3 reasons. Firstly, the *alpha1-tubulin* 3’UTR is only 77bp longer than the *eve* 3’UTR, which is a very minor change relative to the >8kb separating the 5’ and 3’ regulatory elements. Secondly, insertion of much larger MS2 or PP7 arrays (∼1.4 kb) into the *eve* 3’UTR did not disrupt *eve* transcriptional regulation in nc14^20^, indicating that extra sequences are insufficient for ectopic activation of the *eve* minor stripes. Thirdly, the downstream fragment activates minor stripes only during gastrulation^55^, whereas no minor stripe expression was detected at the earlier nc14 stage we studied. We therefore favour the explanation that reduced *eve* mRNA and protein levels alter pair-rule gene expression, allowing ectopic activation of the minor *eve* stripes.

We show that uniform mRNA decay is sufficient to explain the regulation of *eve* stripe 3. We suggest that this relates to the emergence of stripe 3 from an area of low transcriptional activity so, unlike stripes 2 and 4 which refine from broader transcriptional pre-patterns, stripe 3 does not require accelerated mRNA decay at its edges. Our findings highlight how spatially regulated mRNA stability could be an additional strategy for sharpening expression boundaries from broader transcriptional domains (Fig. 6G). This mechanism could function with transcriptional repressors, which are well-established in generating sharp borders in the early *Drosophila* embryo^24,26^ and in vertebrates^56^. Consistent with the rapid pace of early embryonic development, mRNA half-lives in the early embryo are generally shorter than in later tissues, such as the developing nervous system.^7,48^ Therefore, regulation of mRNA decay may provide an extra layer of control for refining precise expression domains over short time scales during embryonic pattern formation.

### Limitations of the study

The modeling approach used here is constrained by data resolution and the challenges of integrating live-imaging and fixed-embryo datasets. We minimized these issues by spatially binning and averaging nuclei, staging fixed embryos using the extent of membrane ingression in nc14, and performing controls for spatial and temporal misalignment. A smFISH-based method for estimating mRNA decay rates has been described.^57^ However, this approach assumes steady-state conditions and is therefore unsuitable for analysing *eve* mRNA stability during stripe refinement, when expression is highly dynamic. Consequently, as described above, our data do not allow us to address spatiotemporal regulation of *eve* mRNA decay. This could be achieved in the future by using spatial transcriptomics to generate time-resolved data that allows paired measurements of transcription and mRNA numbers in the same cells. Although the UTR swap revealed the importance of *eve* mRNA abundance, the ideal experiment would be to alter *eve* mRNA stability at specific embryonic regions, such as stripe boundaries. Future mechanistic and technical advances may enable these *in vivo* perturbations, to extend our *in silico* simulations.

## Resource Availability

### Lead Contact

Requests for further information and resources should be directed to and will be fulfilled by the lead contact, Hilary Ashe

### Materials Availability

All unique/stable reagents generated in this study are available from the lead contact without restriction.

### Data and code availability

- The publicly available MS2 transcription data can be found at: https://cdn.elifesciences.org/articles/61635/elife-61635-supp1-v2.zip. It is programmatically downloaded and processed in the Snakemake pipeline framing the analysis (below).
- The smFISH data can be found on zenodo at https://doi.org/10.5281/zenodo.21068933
- Code for this manuscript is structured into a Snakemake pipeline and is available online at: https://github.com/ManchesterBioinference/spatial_mRNA_decay and at https://doi.org/10.5281/zenodo.21068933.
- Any additional information required to reanalyze the data reported in this paper is available from the lead contact upon request.

## Acknowledgements

We thank Erik Clark for helpful discussions including suggesting that we consider transcript age, Matthew French and Erik Clark for sharing unpublished data, and Tom Pettini for the *eve*, *en* and *wg* smiFISH probes. Thanks to the University of Manchester Bioimaging Facility and Fly Facility, particularly Sanjai Patel for CRISPR injections. This work was funded by Wellcome Trust Investigator and Discovery Awards to H.L.A. and M.R. (204832/Z/16/Z, 204832/B/16/Z, 227415/Z/23/Z) and a Wellcome Trust PhD studentship to J.C.L. (222814/Z/21/Z).

## Author contributions

Conceptualization: L.B., A.P., J.C.L., H.L.A., M.R.; Funding acquisition: H.L.A., M.R.; Investigation: L.B., A.P., J.C.L., C.S., L.F.B.; Software: L.B., J.C.L.; Writing: L.B., A.P., J.C.L., C.S., H.L.A.

## Declaration of generative AI and AI-assisted technologies in the writing process

During the preparation of this work, the authors used Github Copilot, ChatGPT, Gemini, and Microsoft Copilot in order to improve the flow, clarity and grammar of the text. After using this tool/service, the authors reviewed and edited the content as needed and take full responsibility for the content of the published article.

## STAR METHODS

### Biological methods

#### Fly stocks

*Drosophila melanogaster* stocks were maintained at 20°C and raised at 25°C for experiments using standard fly food media (yeast 50 g/L, glucose 78 g/L, maize 72 g/L, agar 8g/L, 10% nipagen in EtOH 27 ml/L, and propionic acid 3ml/L).

The following fly lines were used in this study:

y1w67c23 (BDSC Stock #6599)

y1w*;P{w [+mC] =PTT-GB}me31B[CB05282] (BDSC Stock #51530)

y1sc*v1sev21; P{y[+t7.7] v[+t1.8]=nanos-Cas9.R}attP2 (BDSC Stock #78782)

y1w*; eve-tubulin 3’UTR/ CyO ftz-LacZ (this study)

w[*]; wg[Sp-1]/CyO; ry[506] Sb[1] P{ry[+t7.2]=Delta2-3}99B/TM6B, Tb[+] (BDSC Stock #3629)

#### Genome editing

CRISPR target sites were identified using CRISPR Target Finder (RRID:SCR_023641). Guide plasmids were generated by ligation of 5’phosphorylated annealed oligonucleotides into the BbsI site of pU6-Chi RNA (RRID:Addgene_45946, Gratz et al 2013). Eve homology arms (with mutated PAM sites), replacement coding regions and the *alpha-tubulin84B* 3’UTR (3R:7088584-7088839) swap were inserted into pHD-ScarlessDsRed (DGRC Stock, RRID:DGRC_1364) which had been linearised by PCR. All guide and primer sequences are in Supplementary Table S1.

The donor and guide plasmids were injected into embryos expressing *nos-Cas9* (BDSC Stock #78782) at the University of Manchester Fly Facility. Survivors were crossed to *y^1^w^67c23^* (BDSC Stock #6599) adults to screen for successful CRIPSR events by identifying the presence of DsRed in larvae under a fluorescent stereo microscope. The chromosome carrying *nos-Cas9* was crossed out and the DsRed marker was removed by crossing to the transposase line BDSC #3629 w[*]; wg[Sp-1]/CyO; ry[506] Sb[1] P{ry[+t7.2]=Delta2-3}99B/TM6B, Tb[+] (RRID:BDSC_3629).

#### Embryo fixing and smFISH

For embryo collection, flies were kept in small cages and allowed to lay on apple juice agar plates with yeast paste for 2 hours at 25°C. Plates were then removed and allowed to age for a further 2 hours before dechorionation in 2.5% bleach for 2 minutes and washing with distilled water. Embryos were fixed following the protocol described in ^58^ and stored in methanol at-20°C. Fixed embryos were placed in Wheaton vials (Sigma #Z188700-1PAK) for the smFISH reaction as described.^4^ All smFISH probe sets and stainings were based on the smiFISH protocol.^59^ *eve*, *en* and *wg* smiFISH probes used in this study were a gift from Tom Pettini, the sequences were described previously^60^. *odd*, *ftz, rnt, slp* and *lacZ* smFISH sequences can be found in Supplementary Table S2. The smFISH probes possess a 5’ X, Y or Z-flap sequence^59^, which is recognized by secondary detection probes labelled with Atto 488, Quasar 570 or Quasar 670 fluorophore dyes (LGC Biosearch Technologies). DAPI (500 μg/mL) was added to the third of the final 4 washes of the protocol at a concentration of 1:1000 and embryos were mounted onto slides in ProLong^TM^ Diamond Antifade Mountant (Thermo Fisher Scientific, Cat# P36961) to set overnight before imaging. A Rabbit anti-Eve antibody (1:3000, DSHB, AB_2892600) was used to immunostain embryos in Fig. S5. Embryos for smFISH staining were collected from the following adults: *Me31B-GFP* for Figs. 1-2 and S1-2, *nos-Cas9* for Fig. 3 and S4, and both *nos-Cas9* and *eve-tubulin*/CyO *ftz-LacZ* for Figs. 5-6 and S5-6.

#### Microscopy

A Leica TCS SP8 gSTED confocal microscope was used to acquire the images in Figs. 1-2 and S1-2. For whole-embryo expression domain imaging, a 40x objective with 0.75x zoom was used. For high-resolution imaging of mRNAs and P-bodies a 100x/1.3 HC PI Apo Cs2 objective with 3x line accumulation and 2x zoom was used. The following confocal settings were used for expression domain images: 1 airy unit pinhole, 400 Hz scan speed with bidirectional line scanning, and a format of 4,096 x 4,096 pixels. For high-resolution 100x images the settings were as follows: 0.6 airy unit pinhole, 400 Hz scan speed with bidirectional line scanning, and a format of 2,048 x 2,048 pixels. Laser detection settings were collected as follows: PMT DAPI excitation at 405 nm (collection: 417 to 474 nm); Hybrid Detectors: Alexa Fluor 488 excitation at 490 nm (collection: 498 to 548 nm), Quasar 570 excitation at 548 nm (collection: 558 to 640 nm), and Quasar 670 excitation at 647 nm (collection: 657 to 779 nm) with 1 to 6 ns gating.

All images were collected sequentially, and optical stacks were acquired at system optimized spacing. To allow ageing of the embryos, the membrane was imaged using brightfield at the mid-sagittal plane of the embryo with 40x objective at 0.75x zoom. 1,024 x 1,024 format was used to measure the average length of membrane invagination from at least 5 cells. For all analyses, 3 separate embryos were imaged and quantified as independent replicates. High-resolution smFISH images were deconvolved using Huygens professional deconvolution software by SVI (Scientific Volume Imaging); whole-embryo expression domains images (where single mRNA detection was not carried out) were deconvolved using in-built Imaris deconvolution.

The high-resolution images used for analysis in Fig. 3/Fig. S4 and mRNA counts in Fig. 5B were obtained on a Zeiss LSM 980 with Airyscan2 and a spectral detector 2PMT+32CH-GaASP, in Multiplex SR-8Y mode. Fluorophores were excited using 405/639 nm diode lasers and collected with a Plan-Apochromat 63x/1.40 Oil DIC objective and 3x zoom. Whole embryo images in Figs. 5D, 6, S5 and S6 were acquired similarly but fluorophores were excited using 405/488/561/633 nm diode lasers and collected with a Plan-Apochromat 20x/0.8 M27 objective with 1x zoom. All Airyscan images were captured with bidirectional scanning, 2x averaging, system optimized step size and were processed using Airyscan processing in the Zen blue/black software.

The embryos in Fig. 1 were aged based on 7-9 μm, 10-14 μm or 15-18 μm membrane ingression, but referred to in the figure and text as 8 μm, 12 μm or 16 μm, respectively, for simplicity. All embryo images are shown with anterior to the left.

### Image analysis

#### mRNA and P-body spot detection

Imaris software >9.2.1 (Bitplane, RRID:SCR_007370) was used for nucleus and spot detection. Single mRNAs, TSs and P-bodies were detected separately using the “spots” function, modeling elongation along the z-axis. Detection of nuclei was carried out using the “surfaces” function in Imaris. For quantitation of the number of mRNAs (or TSs) per nucleus, single mRNAs (or TSs) were assigned to the nearest nucleus based on the position of centroids using the sass pipeline ^4^.

#### Eve protein quantification

A 50 μm wide line was drawn down the center of the embryo and the mean intensity of the Eve antibody staining was measured along the AP axis. The mean background intensity from a region of the same area was subtracted. The background subtracted signal was then normalized to the mean background intensity, and peaks from 3 or 4 replicate embryos were manually aligned. The same analysis was repeated for the *eve* smFISH staining.

#### Cuticle preparations

Embryos were aged for 22 hours at 25°C, then dechorionated in 2.5% bleach and washed in 1xPBST. Embryos were washed in 1:1 heptane: methanol for 1 minute before transfer into 1:1 lactic acid: Hoyer’s mounting medium (32% glycerol, 4g/ml chloral hydrate and 0.6g/ml gum acacia), and subsequent mounting onto slides with 22×50mm coverslips. Slides were immediately transferred to 65°C for overnight drying. Cuticles were imaged for quantification using a brightfield tile scan on a Zeiss LSM 980 inverted microscope. Representative darkfield images were taken on a Leica DMR microscope.

### Computational methods

#### P-body colocalization

P-body colocalization analysis across an entire region was carried out as outlined previously using the custom Python code accompanying ^7^ When calculating the P-body colocalization in bins across an image, the positions of colocalized mRNAs, non-colocalized mRNAs and P-bodies in the image were exported. The AP axis was then split into bins corresponding to ∼1 cell width, and the number of colocalized mRNAs (*m_coloc_*) and total number of mRNAs (*m_total_*) within each bin was used to calculate a percentage of mRNAs in P-bodies in each bin across the AP axis. P-body colocalization index (*C_p_*) was calculated as:

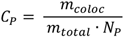

Where *N_P_* is the number of P-bodies.

### Data Acquisition and Preprocessing

#### MS2 Live-Cell Transcription Data

Live-cell MS2 transcription measurements of *even-skipped* (*eve*) expression were obtained from published data.^19^ Supplementary analysis notebooks and CSV files were programmatically downloaded and executed to reproduce the filtered dataset used in the original study, retaining only nuclei exhibiting stable spatial trajectories.

#### Automated Stripe Identification

Stripe positions were identified directly from MS2 fluorescence data. For each nucleus, fluorescence was summarized by summing signal across the final four frames preceding the analysis timepoint (1140, 1160, 1180, and 1200 s). These values capture current stripe positioning while ignoring noise from stripe positional shift with time. Nuclei were binned along the anterior–posterior (AP) axis using bins of width 0.005, and the median fluorescence per bin was computed. The resulting one-dimensional AP fluorescence profile was smoothed using a Savitzky–Golay filter (window length = 17, polynomial order = 3). Stripe centers were detected using prominence-based peak identification (scipy.signal.find_peaks v1.17.0). Stripe boundaries were defined by locating the nearest left and right zero-crossings where the smoothed fluorescence profile fell to zero or below. To ensure symmetric stripe extraction, the smaller half-width was applied on both sides of the peak. Stripes 2, 3, and 4 were extracted for downstream analysis.

#### Spatial Binning of Transcription Traces

Within each stripe, nuclei were partitioned into a 5×5 anterior–posterior × dorsal–ventral (AP×DV) spatial grid. Coordinates were normalized to the observed spatial extent of each stripe.

To reduce discretization artifacts, AP bins overlapped by 50%, while DV bins were non-overlapping. Nuclei contributed to all bins intersecting their spatial coordinates. Mean fluorescence across nuclei was computed at each timepoint, producing 25 positional transcription traces spanning 61 timepoints.

#### Nuclear Density Metrics

To enable later alignment with fixed-cell measurements, embryo-specific nuclear density statistics were computed:

- mean distance to the *k*=4 nearest neighboring nuclei,
- average number of nuclei per spatial bin.

These metrics defined valid density ranges used during smFISH quality control.

#### smFISH Mature mRNA Data

Single-molecule fluorescence *in situ* hybridization (smFISH) measurements of mature *eve* mRNA were collected for stripes 2–4 across two developmental ingression stages (7 µm, 8–9 µm) as explained above. For each stripe, raw Imaris exports were processed using the SASS (https://github.com/TMinchington/sass; github hash 800cb54) pipeline to assign individual transcripts to nuclei.

#### Stripe Centering

Each smFISH image was manually centered along the AP axis when the image was taken and programmatically adjusted as needed. Specifically, transcript positions were binned into 80 AP bins, smoothed using a moving average, and the maximal transcript-density position was identified. When offset from the image center, one AP edge was trimmed until the stripe peak aligned centrally.

#### Density-Based Quality Control and Alignment

To ensure comparability between live MS2 and fixed smFISH datasets, nuclear density constraints derived from MS2 embryos were enforced. Two quality-control checks were applied: (1) Nearest-neighbor density: median *k*-nearest-neighbor distance (*k*=4) required to fall within stripe-specific bounds and (2) Spatial bin occupancy: distribution of nuclei across the 5×5 grid required to match MS2-derived reference ranges.

If smFISH images were too dense, AP edges were iteratively trimmed. If too sparse, the MS2 stripe window was iteratively narrowed and preprocessing recomputed until densities matched. After validation, transcript counts were aggregated into the same 5×5 spatial grid used for transcription data.

### Bayesian Modeling of mRNA Degradation

#### Observation Model

For each embryo, transcription preprocessing yielded 25 spatial transcription traces and smFISH preprocessing yielded a corresponding vector of mature mRNA counts.

Across all models, expected mature mRNA at final timepoint *T* was modelled as a history integral of transcription input weighted by a survival kernel, 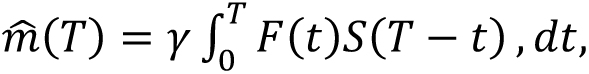, numerically evaluated using the trapezoidal rule. The models differ only in the functional form of the degradation process, which determines the survival kernel. Observed mRNA values were modelled with independent Gaussian noise: 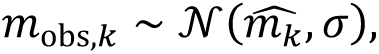, where *k* indexes spatial bins.

All models were inferred using Markov Chain Monte Carlo (MCMC) with the No-U-Turn Sampler (NUTS) implemented in PyMC (v5.27.0) and 4 chains were run with 10,000 iterations each. Parameter estimates were then represented by the mode of samples from the posterior distribution. From parameter estimates of *D*, half-lives were then determined using the following: *t*_1/2_ = *ln*(2)/*D*.

1. *Constant-Degradation Null Model*

As a baseline hypothesis, degradation was assumed to occur at a constant rate independent of spatial position or transcript age. The degradation rate was parameterized by a single scalar (*D_0_*), *D*(*τ*) = *D*_0_. Under constant degradation, transcript survival follows an exponential decay, *S*(*τ*) = exp(−*D*_0_*τ*).

1. *Spatial AP-Dependent Degradation Model*

To test whether degradation varies across embryonic position, degradation rates were allowed to differ between anterior–posterior (AP) spatial bins while remaining constant across dorsal–ventral positions within each AP bin. For AP bin (a), degradation was defined as *D*(*τ*) = *D*_a_. The corresponding survival kernel becomes *S*_a_(τ) = exp(−*D*_a_*τ*), yielding spatially varying but age-independent decay dynamics.

1. *Age-Dependent Degradation Model (Gaussian Random Walk)*

To capture degradation processes that depend on transcript age rather than spatial location, degradation was modeled as a continuous function of molecular age. The logarithm of the degradation profile was parameterized as a Gaussian random walk (GRW), log *D*(*τ_k_*) ∼ GRW, allowing smooth, nonparametric variation in degradation rate across transcript lifetime. The survival probability associated with this age-dependent degradation process is 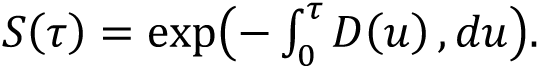.

1. *Biphasic Age-Dependent Degradation via Poly-A Protection*

We further evaluated a mechanistic model in which degradation is regulated by poly-A tail shortening and progressive loss of protection against decay as described^29^ in their modeling of yeast. Poly-A length as a function of transcript age was modeled as *N_A_*(*τ*) = max{0, *N_A_*_,0_ − *rτ*} where (*N_A,0_*) is the initial tail length and (r) is the deadenylation rate. A protection factor was defined as *p*(*τ*) = tanhI*βN_A_*(*τ*)K, which modulates susceptibility to degradation. The resulting age-dependent degradation rate was *D*(*τ*) = *D_protected_* + I1 − *p*(*τ*)K*D_unprotected_*. The implied survival kernel follows 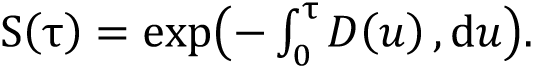.

This model produces biphasic decay behavior in which transcripts are initially protected and become increasingly unstable as poly-A tails shorten.

#### Prior Distributions

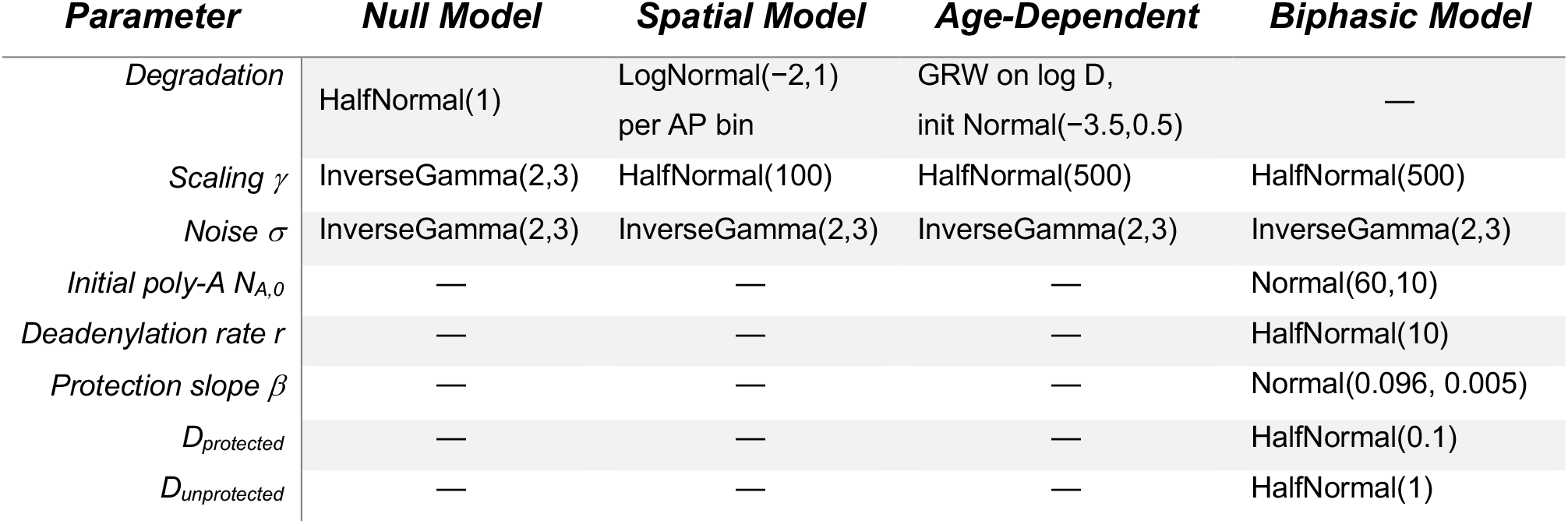

### Model Training Strategy

Each model was fit across three stripes (2-4). Additionally, each model was fit for stripe2 with spatial and temporal shifts to ensure robustness against the uncertainty in the alignment between the MS2 transcription dynamics and smFISH measurements. The temporal shifts were accomplished by imaging embryos at 7um (earlier in time) compared to the 8-9um ingression for the main analysis. The spatial shifts involved shifting the AP axis bins for the MS2 data half a bin’s width to the left or right.

### Model Validation and Comparison

Posterior predictive checks were performed by recomputing expected mRNA levels using inferred parameters and comparing predictions against observed values across all spatial bins. Models were compared using Pareto-smoothed importance sampling leave-one-out cross-validation (PSIS-LOO) implemented in ArviZ (v0.23.0). Observations exhibiting problematic Pareto-*k* diagnostics were refit using exact leave-one-out retraining (reloo). Final model rankings were obtained using expected log predictive density comparisons, identifying the degradation mechanism with the strongest predictive performance.

### Counterfactual Analysis of Uniform mRNA Stability

To assess whether spatial variation in *eve* mRNA stability is required to produce the observed stripe morphology, we performed an *in silico* analysis using the fitted spatial degradation model. Specifically, we generated synthetic mRNA data from the observed transcription traces under scenarios in which mRNA stability was assumed to be uniform across all AP bins. This analysis corresponds to the simulations presented in Fig. 4.

Two uniform half-life scenarios were considered. For the short half-life simulation, a single degradation rate was defined by averaging the degradation rates inferred for the left and right edge AP bins of the fitted spatial model. This represents a scenario in which all positions across the stripe experience the rapid decay observed at the stripe edges. For the long half-life simulation, the AP bin with the lowest inferred degradation rate (highest stability) was identified and its posterior median degradation rate was applied uniformly across all AP bins. This represents a scenario in which all positions across the stripe experience the high stability observed in the stripe center. For both simulations, the inferred transcription scaling factor (*γ*) was held constant. Synthetic mRNA levels were then generated by solving the model using the observed transcription traces and the selected uniform degradation rate. The resulting synthetic mRNA distributions were compared with the experimentally observed mRNA data and with predictions from the fitted spatial model to determine whether uniform mRNA stability is sufficient to reproduce the observed stripe morphology.

### Stripe morphology visualization

Results for stripe 3 indicate spatially varying decay is not essential for stripe boundary refinement, so we investigated the morphology of the stripes’ widths, positions over time, and patterns of emergence to uncover what setsis different about stripe 3 apart in the MS2 data. As described above, stripe ranges across the AP axis for the MS2 data were programmatically identified. Using this method, we calculated the stripe width (max AP position minus min AP position) for each stripe at each time point that it was detected and then visualized it in a line plot. Similarly, the center point for each stripe was calculated ((max AP position + min AP position) / 2) for each stripe at each time point where it was detected. Each stripe’s center position was normalized by the first detected center position, so that each stripe’s center would start at zero to allow for easier comparison between stripes. To visualize stripe emergence, we once again used the same programmatic process used to identify the stripe ranges and positions and applied it to all time points. We then selected the time points best showing the emergence of stripes 2, 3, and 4 as well as the final time point of 1200 seconds for comparison.

### Software Environment

The MCMC workflow constructed with Snakemake was executed using Python 3.11 within a uv-managed environment. Core dependencies included NumPy v2.3.5, pandas v2.3.3, SciPy v1.17.0, scikit-learn v1.8.0, PyMC v5.27.0, ArviZ v0.23.0, and Snakemake v9.14.3. The full pipeline can be reproduced by cloning the repository.

