## Supplementary Figures and Table S1 for "Spatially varying mRNA decay contributes to sharpening the *even-skipped* expression pattern in the *Drosophila* embryo"

A

B

C

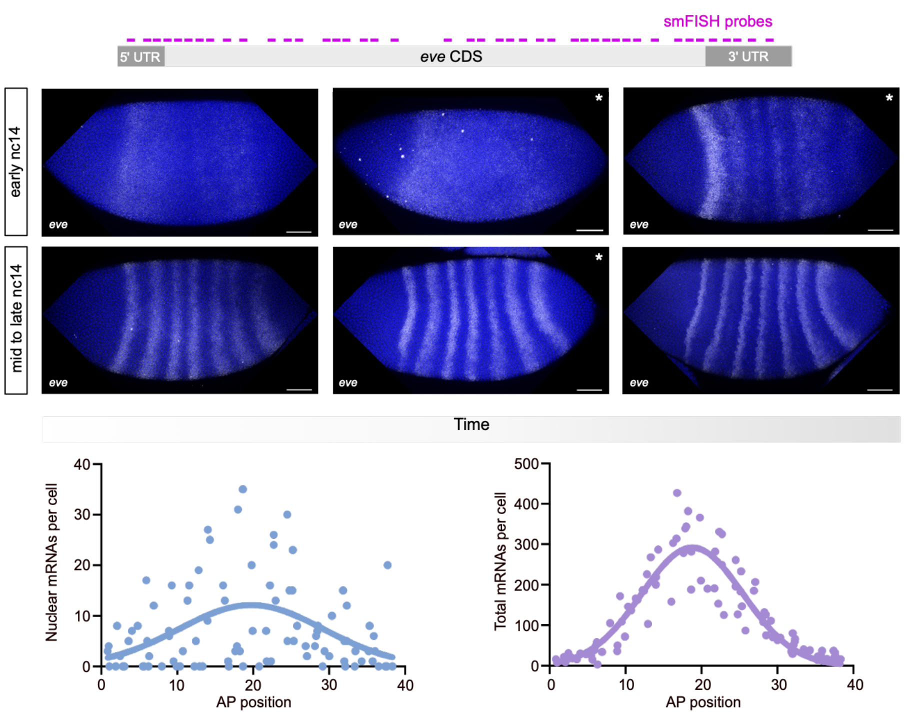


D

nc14 8μm

| AP Bin | 1 | 2 | 3 | 4 | 5 | 6 | 7 | 8 | 9 | 10 | 11 | 12 |
|---|---|---|---|---|---|---|---|---|---|---|---|---|
| mRNAs | 143 | 150 | 499 | 1311 | 2003 | 2062 | 2021 | 1663 | 1022 | 545 | 263 | 197 |
| P-bodies | 720 | 696 | 685 | 643 | 655 | 649 | 682 | 634 | 590 | 634 | 621 | 627 |

nc14 12μm

| AP Bin | 1 | 2 | 3 | 4 | 5 | 6 | 7 | 8 | 9 | 10 | 11 | 12 |
|---|---|---|---|---|---|---|---|---|---|---|---|---|
| mRNAs | 374 | 353 | 349 | 610 | 1262 | 1638 | 1540 | 1305 | 1047 | 768 | 510 | 313 |
| P-bodies | 624 | 538 | 584 | 538 | 491 | 466 | 470 | 471 | 444 | 457 | 455 | 437 |

nc14 16μm

| AP Bin | 1 | 2 | 3 | 4 | 5 | 6 | 7 | 8 | 9 | 10 |
|---|---|---|---|---|---|---|---|---|---|---|
| mRNAs | 235 | 595 | 982 | 1524 | 2505 | 2988 | 2093 | 536 | 57 | 40 |
| P-bodies | 1031 | 1032 | 1036 | 1038 | 931 | 923 | 858 | 891 | 876 | 890 |

E

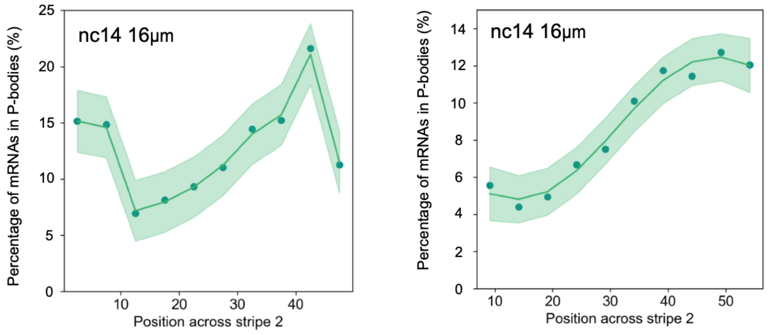


**Figure S1. Analysis of P-body colocalization across *eve* stripe 2, Related to Figure 1.**

(A) Schematic showing the locations of the *eve* smFISH probes across the *eve* mRNA. (B) nc14 embryos stained with *eve* smFISH probes (white) and DAPI (blue). Embryos are ordered in time using the refinement of the *eve* pattern and the extent of membrane ingression in the brightfield channel. The embryos in the panels marked with an asterisk were used to generate the false coloured embryo images shown in Fig. 1A. Scale bars: 50 μm. (C) Graphs show the number of nuclear and total *eve* mRNAs/cell plotted based on AP position, from quantitation of the smFISH image shown in Fig. 1B. (D) Tables show the total numbers of mRNAs and P-bodies per AP bin for each of the embryos shown in Fig 1G. (E) Replicate embryos for P-body colocalization analysis across *eve* stripe 2 at 16 μm membrane ingression. Data over the AP axis are fit with a gaussian process.

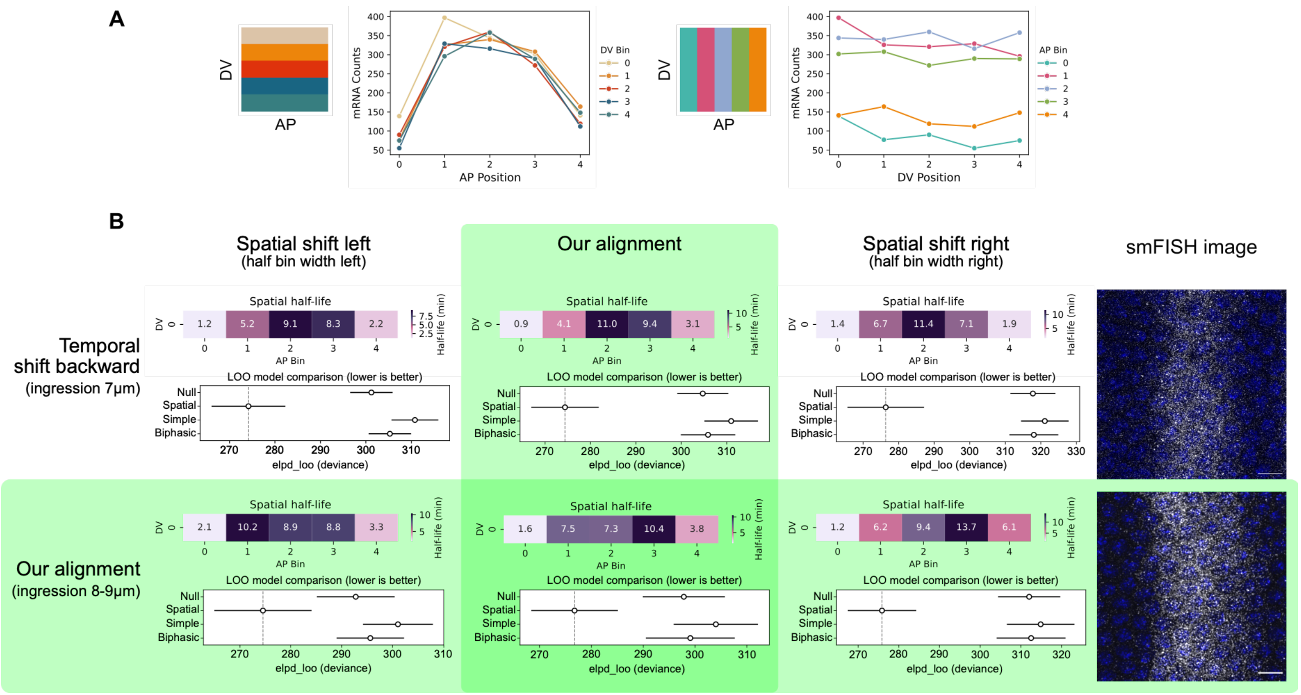


**Figure S2. Modeling results for spatial and temporal shifts using *eve* stripe 2, Related to Figure 2.**
(A) Plots of mRNA counts per bin across the AP and DV axis of the *eve* stripe 2 domain, using data from the embryo shown in Fig. 1B. (B) The spatially defined half-lives and LOO comparison results for stripe 2 and its spatial and temporal shifts to ensure model results are robust to potential misalignments between the MS2 traces and smFISH data. Row 1: temporal shift back in time using an embryo at 7 µm ingression. Row 2: our temporal alignment using an embryo at 8-9 µm ingression to pair with the 1200 timepoint from the MS2 data. Column 1: Spatial shift half a bin's width to the left in the MS2 data. Column 2: our spatial alignment where we programmatically lined up the peak in the MS2 data with the peak in the smFISH data. Column 3: spatial shift half a bin's width to the right in the MS2 data. Column 4: smFISH images of *eve* stripe 2 in the embryos at the two different ingression stages, scale bars: 5 µm. The smFISH data from the embryo in row 2 were used to generate the heatmap shown in Fig. 2B (also shown rescaled in Fig. 4Ai), which was used for the inference in Fig. 2D and Fig. 4Aiii.

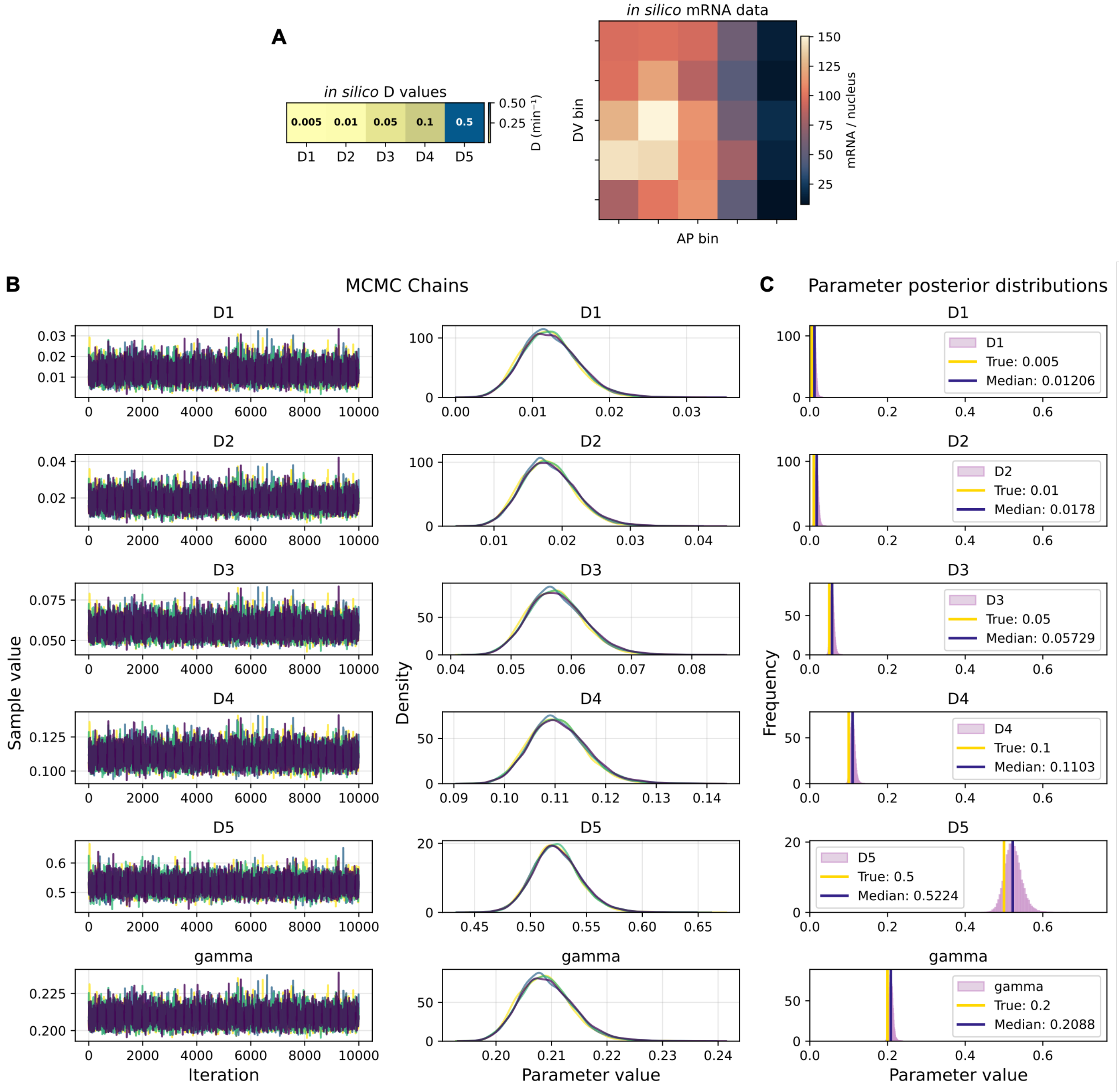


**Figure S3. *In silico* modeling results, Related to Figure 2.**

(A) *In silico* mRNA data, generated using the binned live imaging traces and the generated *D* values displayed here across each AP bin. (B) Markov chains from Monte Carlo sampling of parameters using *in silico* data. Each of the four chains (10,000 iterations each) are shown in different colors. (C) Posterior distributions of parameters from sampling of the *in silico* data. Yellow lines display the true parameter values used to generate the data, and purple lines show the median of the posterior distributions. We note that the model tends to underestimate half-lives (overestimate decay) compared to the true values used to generate the data.

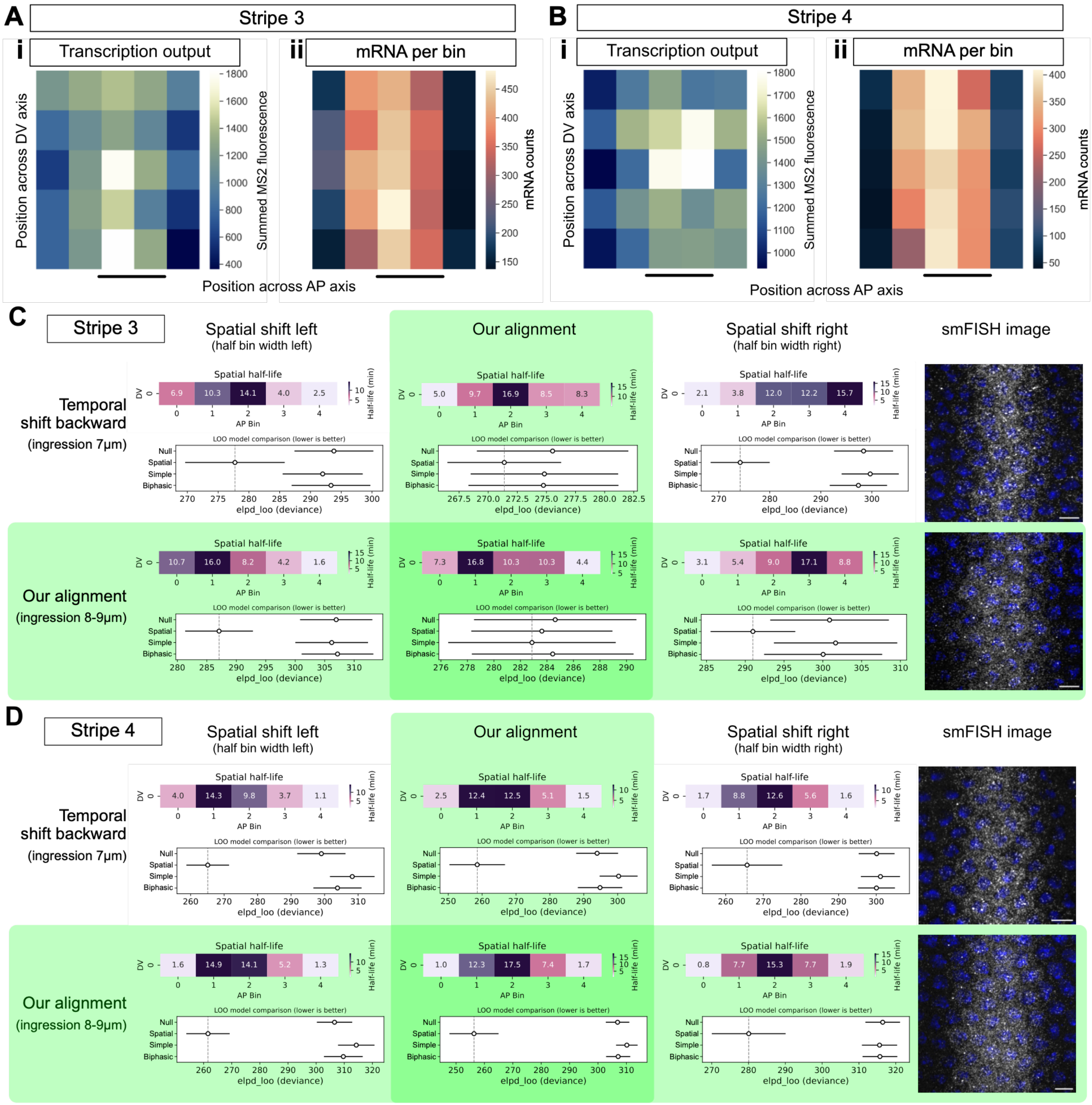


**Figure S4. Modeling results for spatial and temporal shifts using *eve* stripes 3 and 4, Related to Figure 3.** (A-B) Heatmaps of the live imaging *eve*-MS2 data at 20 min into nc14 (Ai, Bi, transcription output) and smFISH data (Aii, Bii, mRNA per cell) at the matched stage for *eve* stripe 3 (A) and stripe 4 (B). The embryo images for stripe 3 and 4 are shown in the lower panels in (C) and (D), respectively, scale bars: 5 μm. The mRNA heatmaps are also shown in Fig. 4Bi, Ci, rescaled to allow easier comparison across the altered half-lives. (C-D) As described for Fig. S2B, but the data are for stripe 3 (C) and 4 (D). The smFISH data from the embryo in the bottom row of (C, D) were used for the inference in Fig. 3A, B and Fig. 4Biii, Ciii.

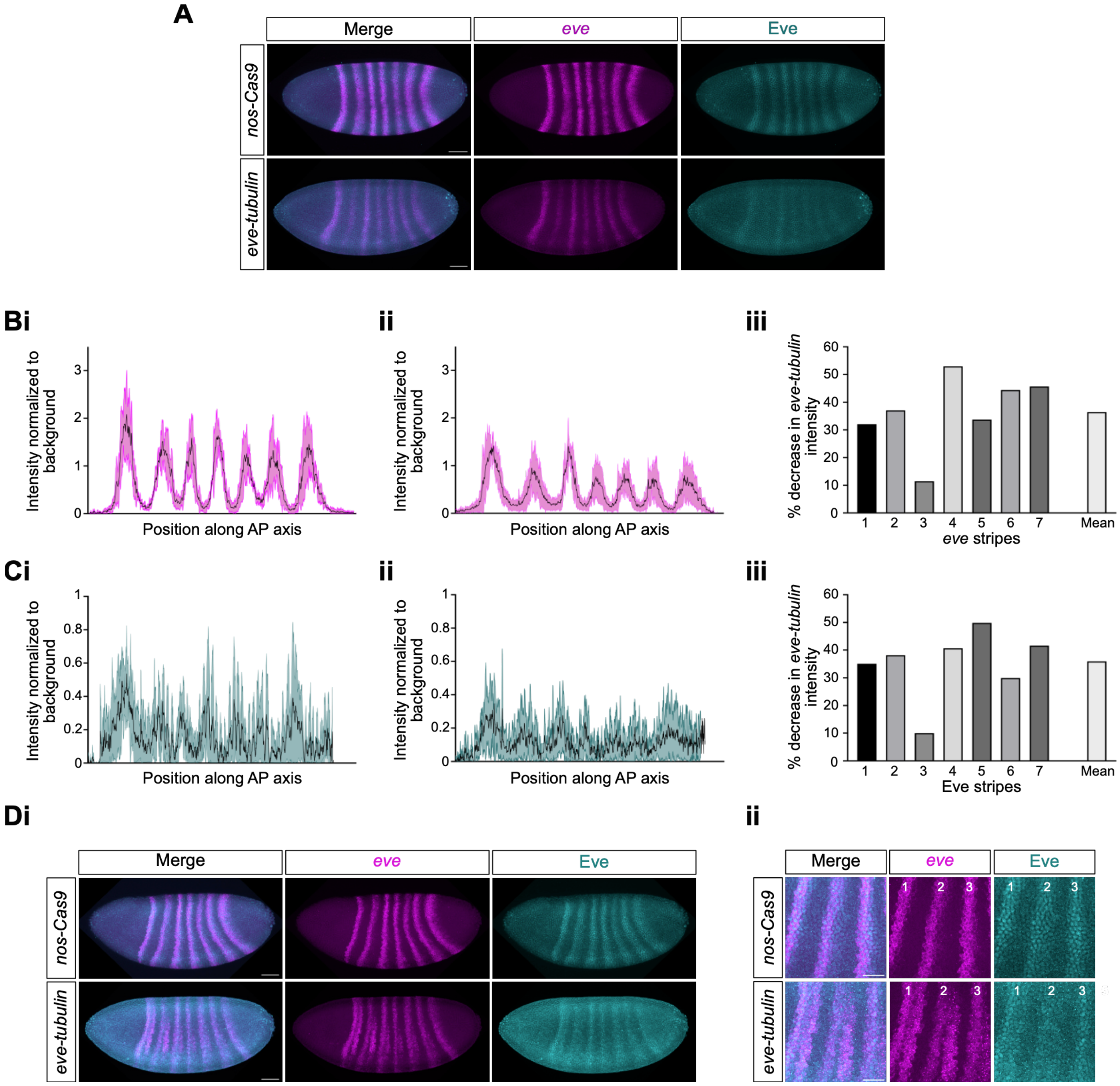


**Figure S5. Reduced *eve* mRNA and protein in *eve-tubulin* embryos, Related to Figure 5.**

(A) Maximum projection images of *nos-Cas9* and *eve-tubulin* embryos at 7-8 µm membrane ingression, stained with *eve* smFISH probes (magenta) and an Eve antibody (cyan). Merged images are shown with single channels for clarity. (Bi, ii) Graphs showing the mean normalised intensity fold change between *eve* mRNA signal and background signal in (i) *nos-Cas9* embryos and (ii) *eve-tubulin* embryos. Data are shown as mean±s.d. value (line and shaded area), n=3 (*nos-Cas9*) and n=4 (*eve-tubulin*). (B iii) Graph shows the percentage decrease in fold-change intensity in *eve-tubulin* embryos compared to *nos-Cas9* controls. The mean is calculated based on the data for all 7 stripes. (Ci-iii) As in (Bi-iii) but for Eve protein intensities. (D) Whole embryo (i) and higher magnification (ii) images of embryos stained as in (A) but at 19-20 µm membrane ingression. The positions of stripes 1-3 are indicated in the higher magnification images. Scale bars: 50 µm, whole embryos, 25 µm, zooms.

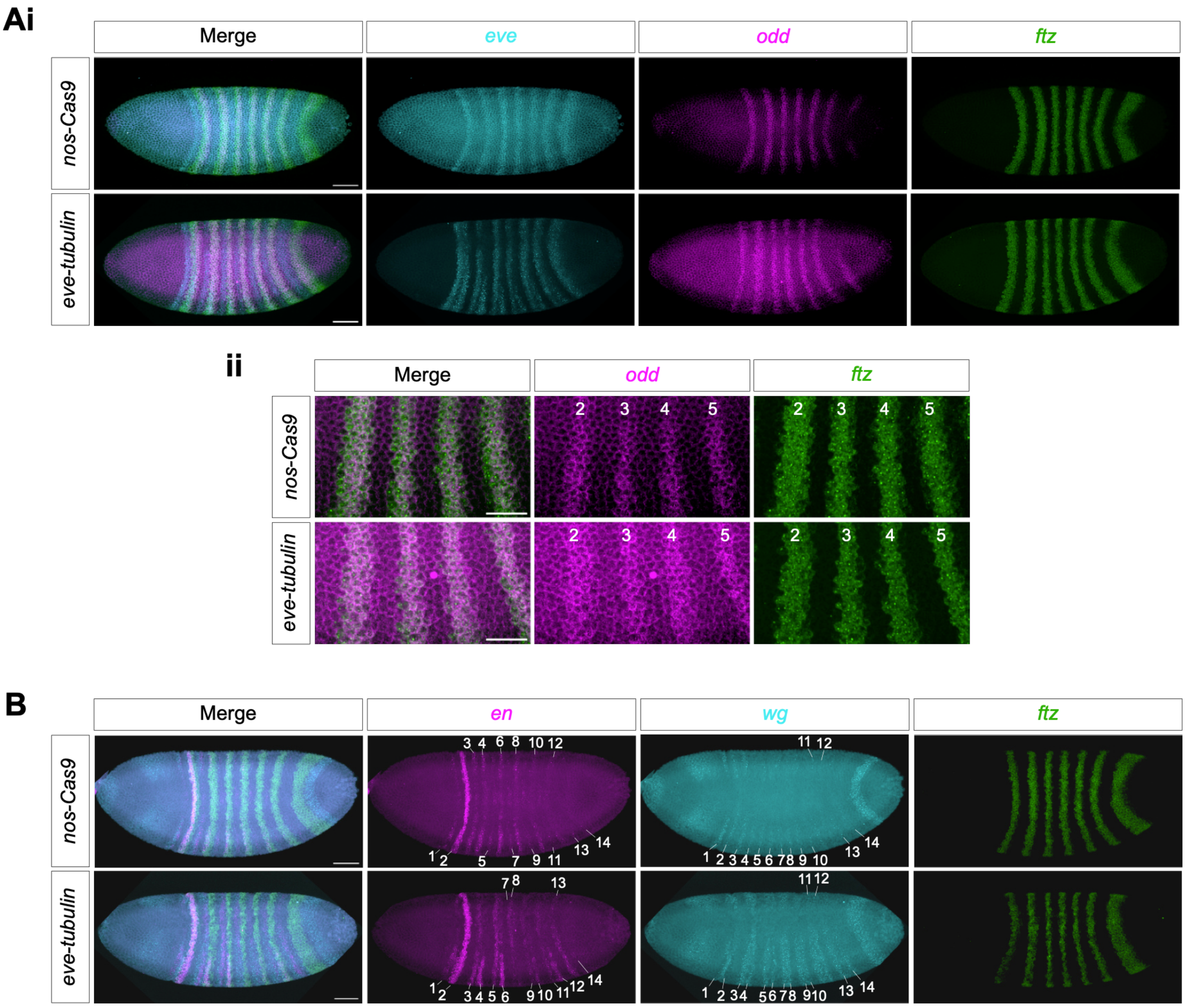


**Figure S6. Disrupted expression of AP patterning genes in *eve-tubulin* embryos, Related to Figure 6.** (A) Maximum projections of smFISH images of *nos-Cas9* and *eve-tubulin* nc14 embryos, with ~25 μm membrane ingression, stained with *eve* (cyan), *odd* (magenta) and *ftz* (green) probes. Whole embryo (Ai) and higher magnification views of stripes 2-4 (ii) are shown as merged images and single channels. (B) Maximum projections of smFISH images of *nos-Cas9* and *eve-tubulin* nc14 embryos, with ~20 μm membrane ingression, stained with *en* (magenta), *wg* (cyan) and *ftz* (green) probes. The 14 *en* and *wg* stripes are labelled. Scale bars: 50 μm, whole embryos, 25 μm, zooms.

**Table S1. PCR primer sequences, Related to STAR Methods.**
PCR primers and CRISPR guide sequences used to generate the *Drosophila melanogaster* line $y^1w^*$; *eve-tubulin*/*CyO ftz-LacZ*.

| Primer Name | Sequence |
|---|---|
| Scarless linearisation 1R | GATCGCAGGTGCTGCCAC |
| Scarless linearisation 1F | TTAACCCTAGAAAGATAATC |
| Eve 1AF | AGGTGGCAGCACCTGCGATCTTCCCAAATGGTTATGGCTGG |
| Eve 1AR | GACAAAGCGGATTAGCAGGACCCGGAGC |
| Eve 1BF | TCCTGCTAATCCGCTTTGTCCCGCCTCG |
| Eve 1BR | GATTATCTTTCTAGGGTTAAACTTACCATTTGGCGAGGGAGG |
| Scarless linearisation 2R | TTAACCCTAGAAAGATAGTCTGCGT |
| Scarless linearisation 2F | GGAAGAGCCGTCGCTCTTCC |
| Eve 2AF | GACTATCTTTCTAGGGTTAAAGATAAAGCCGAGCAAACG |
| Eve 2AR | GGCGTGACGCTTACGCCTCAGTCTTGTAGGG |
| Eve 2BF | TGAGGCGTAAGCGTCACGCCACTTCAAC |
| Eve 2BR | AATACCAGGTACCATTTAGTTCAAATAAACAGTGTGTGTCGTATAATTTTCTTATTTC TGACAACACTGAATCTG |
| Eve 2CF | ACTAAATGGTACCTGGTATTATCTACATATATATC |
| Eve 2CR | GGAAGAGCGACGGCTCTTCCAAAAGTCACTCAACGGGT |
| Guide 1 F: | [P] CTTCGTGTAGATAATACCAGGTACC |
| Guide 1 R: | [P] AAACGGTACCTGGTATTATCTACAC |
| Guide 2 F: | [P] CTTCGAACGAGGCGGGACAAAGCGG |
| Guide 2 R: | [P] AAACCCGCTTTGTCCCGCCTCGTTC |
